# The PSInet Plant Water Potential Database: advancing new perspectives on plant water status, traits, and hydraulic processes

**DOI:** 10.64898/2026.09.17.752364

**Authors:** Jessica Guo, Ana Maria Restrepo Acevedo, Marvin Browne, Daniel M. Johnson, Katherine A. McCulloh, Jesse B. Nippert, Rafael Poyatos, Steven A. Kannenberg, Daniel P. Beverly, Arthur Endsley, Andrew F. Feldman, Alexandra G. Konings, Yanlan Liu, Jordi Martínez-Vilalta, William M. Hammond, Kevin R. Hultine, Lauren E.L. Lowman, Jeffrey S. Dukes, Julia K. Green, Lawren Sack, Nidhi Vinod, Jie Hu, Jean Allen, Caleb E. Adams, Henry D. Adams, Lucette L.A. Adet, Anthony Ambrose, Leander D.L. Anderegg, William R.L. Anderegg, Luiza Maria T. Aparecido, Ismael Aranda, Eleinis Ávila-Lovera, Eduardo Arcoverde de Mattos, Kinzie C. Bailey, Dennis D. Baldocchi, Maoya Bassiouni, Enric Batllori, Berit E. Batterton, Taylor Baugh, Wendy L. Baxter, Michael C. Benson, Carl J. Bernacchi, Daniel Berveiller, Chris J. Blackman, Davis E. Blasini, Benjamin W. Blonder, Gil Bohrer, Damien Bonal, Ben Bond-Lamberty, Indra Boving, David D. Breshears, Mario Bretfeld, Craig R. Brodersen, Timothy J. Brodribb, Callum Bryant, Sandra J. Bucci, Erika R. Bucior, Maria C. Caldeira, Otávio C. Campoe, Laura V. Cano-Arboleda, Luisina M. Carbonell-Silletta, Amanda A. Cardoso, Raiza J. Castillo Argaez, David Chaparro, Brendan Choat, Deborah Corso, Sabrina Coste, Danielle Creek, Alex E. Crookshanks, Marcella Cross, Jessica L. Cruz, Teresa S. David, Kenneth J. Davidson, Renaud Decarsin, María Del Rey-Granado, Nicolas Delpierre, Sylvain Delzon, Renata M. Diaz, Lee T. Dickman, Frederic C. Do, Jean-Christophe Domec, Tingfa Dong, Avery W. Driscoll, Stefanie Dumberger, Cleiton B. Eller, Manuel Esperon-Rodriguez, Brent E. Ewers, Xue Feng, Victor Flo, Dulce Flores-Rentería, Patrick Fonti, Alicia Forner-Sales, Rafael S. Freitas, Núria Garcia-Forner, Jess Gersony, Mana Gharun, Bruno O. Gimenez, Teresa E. Gimeno, Gregory R. Goldsmith, Ignacio Gonzalez-Fernandez, Sybil G. Gotsch, Anne Griebel, Charlotte Grossiord, Huade Guan, Joannès Guillemot, Simon Haberstroh, Christina A. Hackmann, Martina Hájíčková, Erik P. Hamerlynck, Beatrice L. Harrison Day, Virginia Hernandez-Santana, Natan Holtzman, Louise A. Hosburgh, Jianbei Huang, Brett A. Huggett, Nicole M. Hughes, Valeriy Ivanov, Hervé Jactel, Jaideep Joshi, Ansgar Kahmen, Lee A. Kalcsits, Medelin E. Kant, Lucy P. Kerhoulas, Tamir Klein, Anne Klosterhalfen, John F. Knowles, George Koch, Dan F. Koepke, Natalia Kowalska, Gabriel T. LaHue, Julien Lamour, Victor Lechuga, Sung-Ching Lee, Pedro A.M. Leite, Christoph Leuschner, Jean-Marc Limousin, Juan Carlos Linares, Raquel Lobo-do-Vale, Rosana López, Ana López-Ballesteros, Guillermo Lopez Castro, Cate Macinnis-Ng, Nicolas Martin-StPaul, Ashley M. Matheny, Ilaine S. Matos, Marie Matoušková, Nate McDowell, J. Patrick Megonigal, Patrick Meir, Maurizio Mencuccini, Sean T. Michaletz, Jose Carlos Miranda, Justine E.C. Missik, Teresa Morán-López, Luna Morcillo, Kendalynn A. Morris, Malini Muthu Karpagam, Rachael H. Nolan, Kiona Ogle, Rafael S. Oliveira, Anna Jiselle S. Ongjoco, Elsa M. Ordway, Sharath S. Paligi, Sari Palmroth, Mohan Pandey, Robert E. Pangle, Drew M.P. Peltier, Josep Penuelas, Jennifer M.R. Peters, Richard L. Peters, Clara A. Pinto, R. Suzuky Pinto, Alexandria L. Pivovaroff, William T. Pockman, Quentin Ponette, Daniel L. Potts, Estela Quintero-Vallejo, Mukund P. Rao, Boris Rewald, Trent W. Robinett, Jesús Rodríguez-Calcerrada, Ido Rog, Alistair Rogers, Víctor Rolo, Bruno H.P. Rosado, Lucy Rowland, Raquel Ruiz-Checa, Yann Salmon, Roberto L. Salomón, Adriana Sanchez, Leticia Sandoval, Louis S. Santiago, Cristina Santos da Silva, Gerard Sapes, Fabio Scarpa, Konstantin Schellenberg, Fabian G. Scholz, Aabhas Senapati, Shawn P. Serbin, Hernán Serrano-León, Ankit Shekhar, Ladislav Sigut, Guillaume Simioni, Milos Simovic, William K. Smith, Kevin Soland, Gustavo C. Spanner, Clément Stahl, Zhongbo Su, Jennifer J. Swanson, Yidong Tong, Tonantzin Tarin, Matthew W. Tomazewski, Amy M. Trowbridge, Daphna Uni, Josef Urban, Coral del Mar Valle Rodríguez, Joseph G. Verfaillie, Alberto Vilagrosa, Lorenz Walthert, Lixin Wang, Wenzhi Wang, Nicholas D. Ward, Jay W. Wason, Christiane Werner, Adam G. West, Jason B. West, Virginia G. Williamson, Stephanie J. Wilson, Jeffrey D. Wood, Evan Wyers, Enrico A. Yepez, Yijian Zeng, Yao Zhang, Zhechen Zhang, Camille Ziegler, Kimberly A. Novick

## Abstract

Water potential (Ψ) gradients drive water flow within and between soils and plants, and the internal plant Ψ controls a wide range of physiological processes including photosynthesis, growth, and mortality. Notwithstanding this clear relevance for many critical aspects of ecosystem function, Ψ data have historically been relatively inaccessible and unnetworked. The absence of a centralized repository for plant Ψ time series limits our ability to integrate a wealth of ecophysiological information from other networks and from remote sensing. Closing this gap is necessary to address unresolved questions about plant responses to drought and heat stress, and to make confident predictions about plant and ecosystem function in a warming world. Here, we introduce the PSInet database – a global collection of plant water potential time series from 285 datasets representing 523 species. We present the workflow that guided database development and evaluate its key features. Through a series of preliminary analyses, we then highlight the potential of the PSInet database for applications including: a) advancing plant water use strategy frameworks; b) disentangling the impacts of soil versus atmospheric drought stress; c) assessing the long-held assumption of pre-dawn equilibration of ecosystem water potential; d) understanding the risk of drought-driven mortality; and e) benchmarking remote-sensing data products and land-surface models.

## 1. Introduction

Of all the ways that climate change affects plants, feedbacks linked to increasing drought and heat stress are among the most harmful and the most difficult to characterize. Massive tree die-offs are increasingly common (Hammond et al. 2022; McDowell et al. 2022), and plant carbon uptake and growth are being altered by climatic stressors in ways that have profound impacts on the provisioning of food and timber supply, biodiversity, and the land carbon sink (Green et al. 2019; Liu et al. 2020; Novick et al. 2024; Peters et al. 2025). Anticipating and mitigating these deleterious impacts requires mechanistic and predictive modeling frameworks. However, despite many recent theoretical and conceptual advances (e.g., Mencuccini et al. 2019; Potkay et al. 2025; Sperry et al. 2016; Wolf et al. 2016), key questions about plant water use strategies are unresolved (Kannenberg et al. 2022), and the development and benchmarking of models that robustly predict plant function in a warmer world remains difficult (Corak et al. 2024; Green et al. 2024).

Many gaps in our understanding of plant responses to drought and heat stress can be traced to a historic lack of accessible data on plant water potential (hereafter Ψ, Kannenberg et al. 2022; Konings et al. 2021; Novick et al. 2022; Restrepo Acevedo et al. 2024). The total Ψ describes the free energy of water, and it can be represented as the sum of three components. The first is **the pressure potential,** which integrates the hydrostatic and matric components that tend to dominate plant Ψ (Boyer et al. 1976; Genard et al. 2001). In unwilted plant cells, the pressure potential is positive and often called turgor pressure. In xylem, it is negative, representing tension forces (de Swaef et al. 2022; Peters et al. 2021; Pockman et al. 1995). The second is **the osmotic potential**, which is the potential due to solutes. It is a negative component of water potential within plant cells but often assumed to be zero in the xylem (Rodriguez-Dominguez et al. 2022). The third is **the gravitational potential,** which reduces the total Ψ by 0.01 MPa per meter of height above the ground and is an important component of Ψ in tall woody plants (Koch et al. 2004).

Gradients in Ψ along the soil-plant-atmosphere continuum are the driving force of ecosystem water flows (e.g., Sperry et al. 1998; Tyree & Zimmerman 2002), with water flowing from higher to lower water potentials. Moreover, the magnitude of Ψ directly controls plant growth and mortality during periods of hydrologic stress. Specifically, the Ψ of leaves and stems become more negative during drought and heat waves, reflecting the combined pressures of declining soil water, depletion of the storage tissues, increasing atmospheric demand for water vapor, and a greater need for transpirational cooling (de Kauwe et al. 2019; Martínez-Vilalta et al. 2017; Mencuccini et al. 2024). If leaf Ψ becomes too low, it can lead to leaf wilt and potentially cause lethal embolisms that propagate through the xylem (Tyree & Sperry 2002). As a result, plant stomata tend to close during drought to buffer against these dangerous declines in Ψ (Jarvis 1976). However, stomatal closure leads to reductions in photosynthesis, and even relatively mild water potentials can lead to strong growth cessation (Muller et al. 2011). Moreover, stomatal regulation is imperfect, and incomplete stomatal closure and cuticular transpiration can cause additional water losses from plants and soil (Duursma et al. 2019).

Other traits -- including the turgor loss point in leaves, xylem vulnerability to embolism, and the quantum efficiency of PSII photochemistry -- are important for determining whether damage to cells and xylem networks actually occurs at a given value of Ψ (Bartlett et al. 2012). An extensive body of literature explores how these traits are coordinated with environmental factors to determine plant water use strategies across species, growth habitats, and over time (Kannenberg et al. 2022; McCulloh et al. 2019; Rowland et al. 2023). This information is useful not only for advancing basic science but also for testing and improving land-surface and hydrologic models (e.g, Jimenez-Rodriguez et al. 2024). However, historically, a lack of accessible data describing plant Ψ has impeded the validation and advancement of plant water use frameworks (Kannenberg et al. 2022).

This data scarcity does not reflect a failure to measure Ψ *per* se. Indeed, for decades, observations of leaf and stem *Ψ* have been a cornerstone of ecophysiological research. They are most often made with a “pressure chamber” (Scholander et al. 1965), which has the advantage of being easy to operate under a wide range of field conditions. However, because the approach is laborious and destructive (Rodriguez-Dominguez et al. 2022), pressure chamber data are discrete, often made at timescales of weeks or longer. In-situ, continuous measurement of plant Ψ is possible from psychrometers and microtensiometers (Guo et al. 2020; Lasko et al. 2022; Wang et al. 2014), though these instruments can be sensitive to fluctuations in moisture and temperature (Restrepo-Acevedo et al. 2026). It is also possible to infer Ψ from the pressure volume (or PV) curve, which relates Ψ to the plant volumetric water content (Tyree & Hammel 1972). Recently, there have been significant advances in techniques for continuous monitoring of plant water content *in situ* using automated dendrometers (Bourbia et al. 2025; Peters et al. 2025), time-domain reflectometry (Restrepo-Acevedo et al. 2020), and remotely using tower- or satellite-based microwave measurements (Konings et al. 2021).

Rather, the water potential data gap arises from the fact that, until now, there has been no centralized, accessible database of plant Ψ timeseries, which is particularly striking given the recent proliferation of networks and databases that aggregate and openly share data describing other aspects of plant function. These include: flux tower networks that monitor ecosystem carbon, water, and energy exchanges (Delwiche et al. 2024); the SAPFLUXNET global database of tree water use (Poyatos et al. 2021); the Xylem Functional Traits (XFT) database, which provides species-level information on many traits that determine tree mortality risk (Choat et al. 2012, https://xylemfunctionaltraits.org/); the Regional Ecosystem Soil Hydraulics Project (ReESH), which provides soil hydraulic properties for ecological research sites (Crookshanks et al. *under review*), and nascent networks of automated dendrometer timeseries data (e.g. TreeNet, Zweifel et al. 2021, and www.globaldendro.org). At the same time, satellite and reanalysis data products, including soil moisture and vegetation water content, present new opportunities to scale field-based plant hydraulics knowledge across the globe (Feldman et al. 2021; Haynes et al. 2026; Konings et al. 2021; Liu et al. 2021). However, in the absence of similarly accessible information about Ψ, we are limited in our capacity to synthesize information across sites and with remote-sensing and modeling platforms.

Here, we introduce the PSInet database – a new, global, open-access database of plant water potential time series that aggregates both pressure chamber and automated measurements of plant Ψ. The database is a product of the PSInet research coordination network (Restrepo-Acevedo et al. 2024), which is a global initiative supported by the U.S. National Science Foundation (NSF) to promote the collection, sharing, and synthesis of Ψ data (https://psinetrcn.github.io/). A key feature of this database is its focus on enabling integration with data from other environmental observation networks and remote-sensing platforms, and with land surface modeling frameworks. We describe the workflow by which the database was created and details on how the database can be accessed. Then, we present preliminary analyses that can motivate effective end-use of the database, alone and in combination with data from other networks. These applications include conceptual frameworks for plant water use strategies, the disentanglement of soil versus atmospheric drought impacts, more robust characterization of plant mortality risk, and the advancement of remote sensing products and process-based models.

## 2. Database Creation and Access

The initiative began in 2022, when a steering committee was formed to envision a database (Fig. 1) comprising plant Ψ timeseries, associated metadata, and optional soil and meteorological data. Datasets were solicited from 2024 until early 2026, at which point submissions were closed. The database described here is thus ‘static,’ with the possibility of re-opening the database in the future contingent on funding and other resource availability. Plant Ψ time series were defined as multiple measurements on the same individual or group of individuals, and we collected metadata describing site, treatment, plot, and plant characteristics, including availability of data from other databases. No minimum temporal resolutions or durations were enforced in order to maximize community data contributions. We acknowledge that not all datasets are suitable for all proposed applications, and database end users are encouraged to query the database with filters suitable for their specific application. Harmonizing metadata across experimental designs was challenging: some studies intensively and repeatedly measured plant Ψ at a single site while others sampled plant Ψ across multiple sites, species, or individuals at lower frequency.

**Fig 1.**
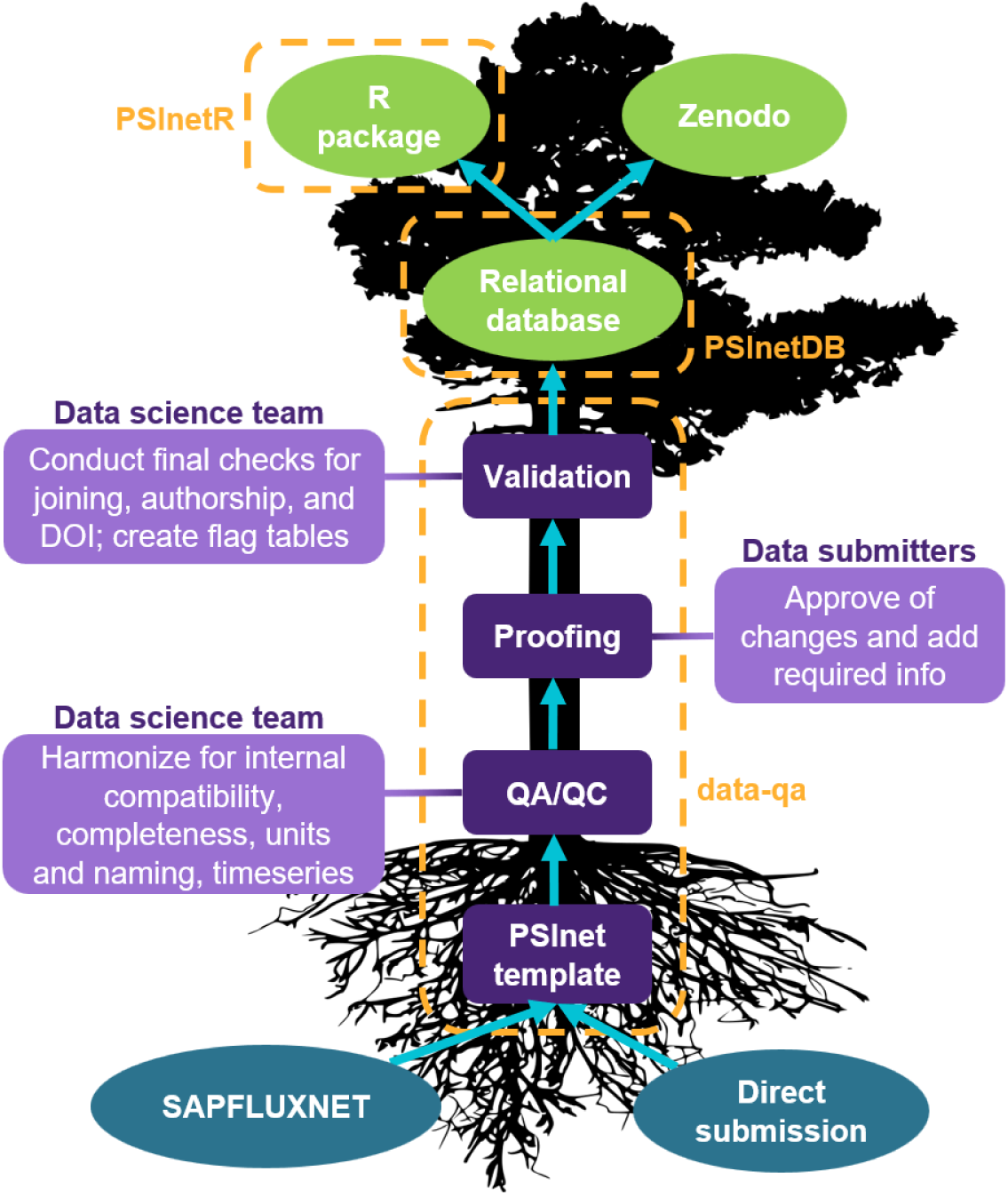
Diagram depicting the PSInet workflow for ingesting data and database creation. Data were submitted from two sources (SAPFLUXNET or direct submission) in the PSInet template format. After initial QA/QC, data were returned to data submitters for proofing. Validation of the returned proofs yielded the raw material for creating the relational database. Database tables are additionally available as rectangular .csvs on Zenodo and as an R package with custom joining functions that output ready-to-analyze products. Dashed boxes and labels indicate the associated GitHub repository.

Data providers shared data using a common Excel template that accommodated a wide range of experimental designs, attributed treatments at the plot or individual level, and nested individuals within plots and sites. Naturally-occurring events were classified as treatments if they varied in space and on distinct individuals (i.e., crown dieback, mistletoe infection, disturbance level) but not if varying in time and affecting all individuals (i.e., heat wave). We defined individuals as belonging to only a single treatment in order to standardize database structure and table relationships. Multi-site studies had separate templates for each location, which were subsequently linked during the QA/QC process. A sensor metadata table was added to describe the sensors associated with automated plant Ψ; additional flexibility was necessary because multiple sensors could be instrumented on the same plant and because the same sensor could be rotated among plants at different times.

Completed templates underwent a human-led, script-based, and reproducible QA/QC process that verified completeness, unit and naming consistency, and cross-compatibility among tables. Expert judgement was used to determine what corrections were needed, if any, and the resulting data were returned to data contributors in proof format for further correction and approval. The resubmitted proofs underwent a final set of checks before all tables were saved as flat .*csv* files. Species names were checked against multiple taxonomic databases using the package ‘taxize’ (Chamberlain et al. 2020). Additional tables for binary quality flags were produced for all time series (chamber and automated Ψ, soil moisture, meteorological variables) using fixed range requirements (Table S1). Range requirements were adapted from the SAPFLUXNET database (Poyatos et al. 2021), with relative humidity and vapor pressure deficit ranges lightly modified to account for the greater number of arid sites in PSInet. Files were stored on Google Drive, processed using R, and version controlled with git and GitHub.

Best practices for the collection and quality control of automated water potential data are less well established (but see Restrepo-Acevedo et al. 2026), with considerable variability in protocols for sensor operation, placement, calibration, and data screening from one study to the next. For this reason, we exercised minimal QA/QC of the automated water potential data, refrained from gapfiling, and used only range checks to produce quality flags. We note that some data contributors used their own QA/QC prior to submission, while others shared raw sensor data. Users are strongly encouraged to exercise their own scientific judgement and practice custom QA/QC on all automated timeseries variables. We hope that the PSInet database can spur the creation of community-developed standards for the QA/QC of automated water potential timeseries.

Following quality control, individual submissions were aggregated into a relational database using the DuckDB format. The database consists of 18 tables, with the ‘study_site’ table containing the primary key of ‘dataset_name’, the unique identifier of each submission that links with all other tables. Users can access the database via the R package ‘PSInetR’, which delivers the psinet.duckdb file locally. From there, users can query the database with standard ‘dplyr’ functions (Wickham et al. 2026a); the backend package ‘dbplyr’ provides translation into SQL (Wickham et al. 2026b). Custom functions within ‘PSInetR’ join the metadata tables with chamber plant Ψ, automated plant Ψ, soil moisture timeseries, and meteorological timeseries as rectangular dataframes. The DuckDB format is language agnostic; non-R users can access the database via the command line, Python, or third-party GUI interfaces.

The PSInet database will be available under a CC BY 4.0 license following a one-year embargo period which ends on October 1, 2027. Following the PSInet data policy, data contributors will have advanced access to the database during the embargo period, after which point, the database will become open to all potential end-users provided they follow the proper attribution protocol (https://psinetrcn.github.io/data_terms.html). Specifically, future end-users of the database must cite this paper in the main text of any journal article that makes use of PSInet data, and include the following text in the acknowledgments “*Funding for the creation of the PSInet database was provided by the US National Science Foundation administered by the Division of Integrative Organismal Biology via a Research Coordination Grant (#2243900).”* Additionally, if five or fewer datasets are utilized, end-users must also provide a citation for each unique dataset (if available) in the main text of journal articles. In this case, database end-users are strongly encouraged to contact the dataset contributors to invite their collaboration on the paper, though this cannot be strictly enforced.

## 3. Key features of the PSInet database

A total of 285 water potential time series representing over 625,000 observations from six continents were submitted to PSInet (Fig. 2a, Tables S2 and S3). The vast majority of the datasets (n = 273) were collected in the field, with a handful of data from greenhouse studies (n = 12). In some cases, multiple datasets originated from the same study, but differed in terms of location, instrumentation (chamber versus automated), or some other fundamental aspect. For consistency, we describe each dataset as the unit of submission, while noting that datasets originating from the same study may not always be independent in space or time. Pressure chamber time series dominated the submissions (280 time series representing >99,000 observations, Fig. S1). The submission of automated water potential data was more limited (n = 16), but these data provide substantially more observations (> 528,000, Fig. S1). Eleven datasets included both automated and chamber timeseries. Out of 285 datasets, 64 are co-located with eddy covariance (EC) towers, 39 with SAPFLUXNET data, and 7 with both networks. Nine out of 16 automated datasets are co-located with EC towers. For a summary of dataset characteristics, see Tables S2 and S3.

**Fig. 2:**
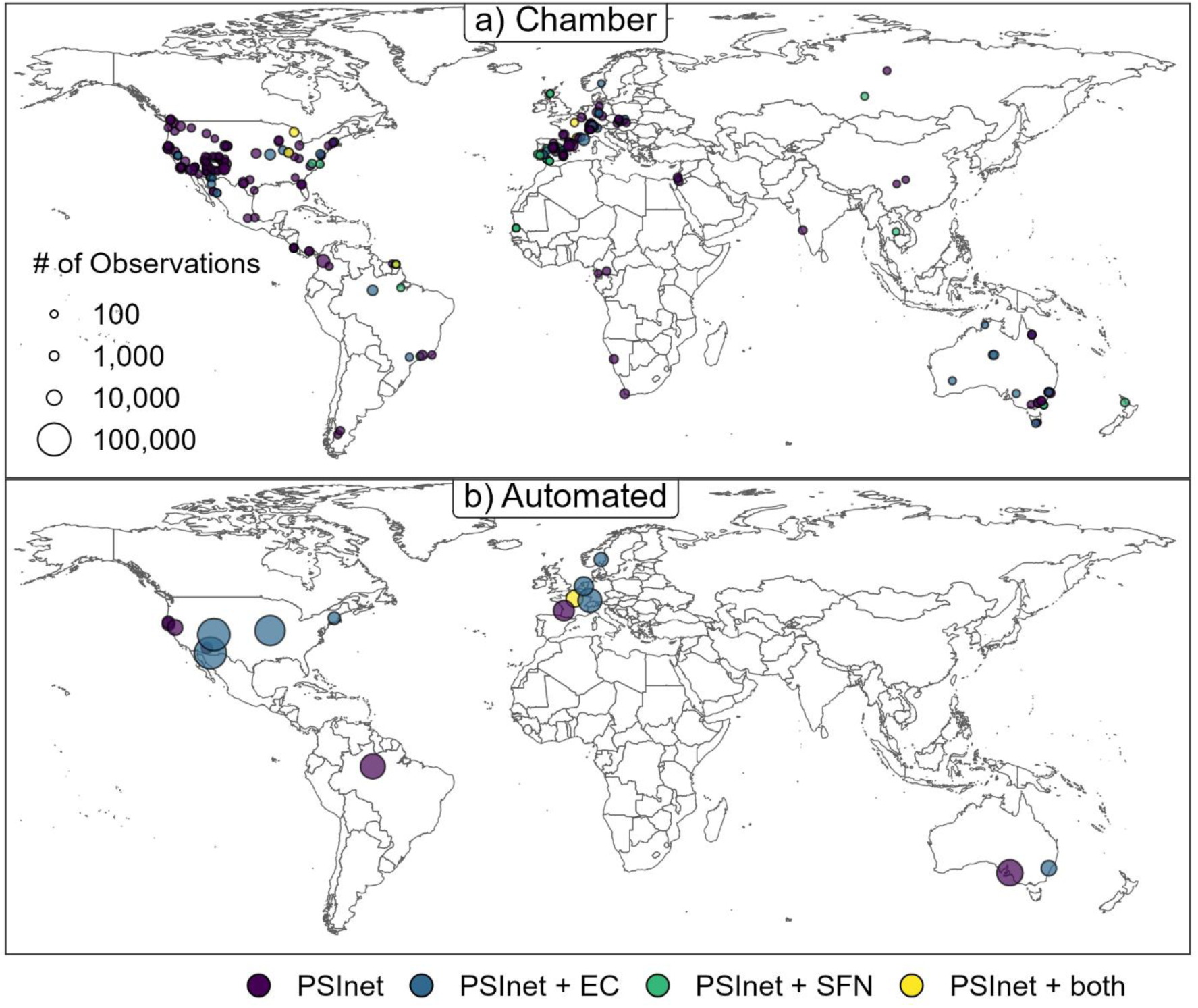
Spatial distribution of PSInet pressure chamber (a) and automated (b) measurements. Size of points are scaled by number of observations; colors represent co-location with eddy covariance towers (EC), SAPFLUXNET data (SFN), or both.

The most well-represented Whittaker biomes (based on mean annual precipitation and mean annual temperature from WorldClim) in the database (Fig. S2) include arid woodlands and shrublands (n = 106), temperate seasonal forests (n = 69), subtropical deserts (n = 36), temperate grasslands/deserts (n = 27), and tropical seasonal forests and savannas (n = 22). Temperate rain forests (n = 14) were reasonably well represented, but only a handful of submissions originated from tropical rain forest, boreal forests, and tundra (n = 6, 4, and 1 respectively). The lack of representation from tropical biomes is a well-documented and persistent shortcoming that cuts across many environmental observation networks (de Aguiar-Campos et al. 2025; Villareal & Vargas et al. 2021). Increasing representation from the tropics and other underrepresented biomes should continue to be an emphasis of future environmental network development.

The database contains records from more than 523 unique plant species from 105 families (Fig. 3a); in 12 instances, plants were only identified to the genus level. The vast majority of species were angiosperms (n = 470), of which most were eudicots (n = 402). The rest were split between magnoliids (n = 34) and monocots (n = 34). Gymnosperms were well-represented (n = 50), but very few pteridophytes are represented in the database (n = 3). Species were then classified into nine simple plant functional types using the TRY Categorical Traits Dataset (Kattge et al. 2011, Kattge et al. 2012) and supplemented by author-provided traits and additional online searching. Woody PFTs, comprising trees and shrubs, are extremely well-represented in the database (n = 459), whereas only 40 forb/herb species and 24 grasses are present. For chamber observations, the top ten most represented species were tree species (Fig. 3b), and *Pinus edulis* and *Quercus ilex* were represented by the greatest number of datasets (n = 18 and 20, respectively). The top ten species by number of automated observations (Fig. 3c) were dominated by plants from more arid climates where psychrometers work particularly well, including *Juniperus monosperma* and *Larrea tridentata*.

**Fig. 3:**
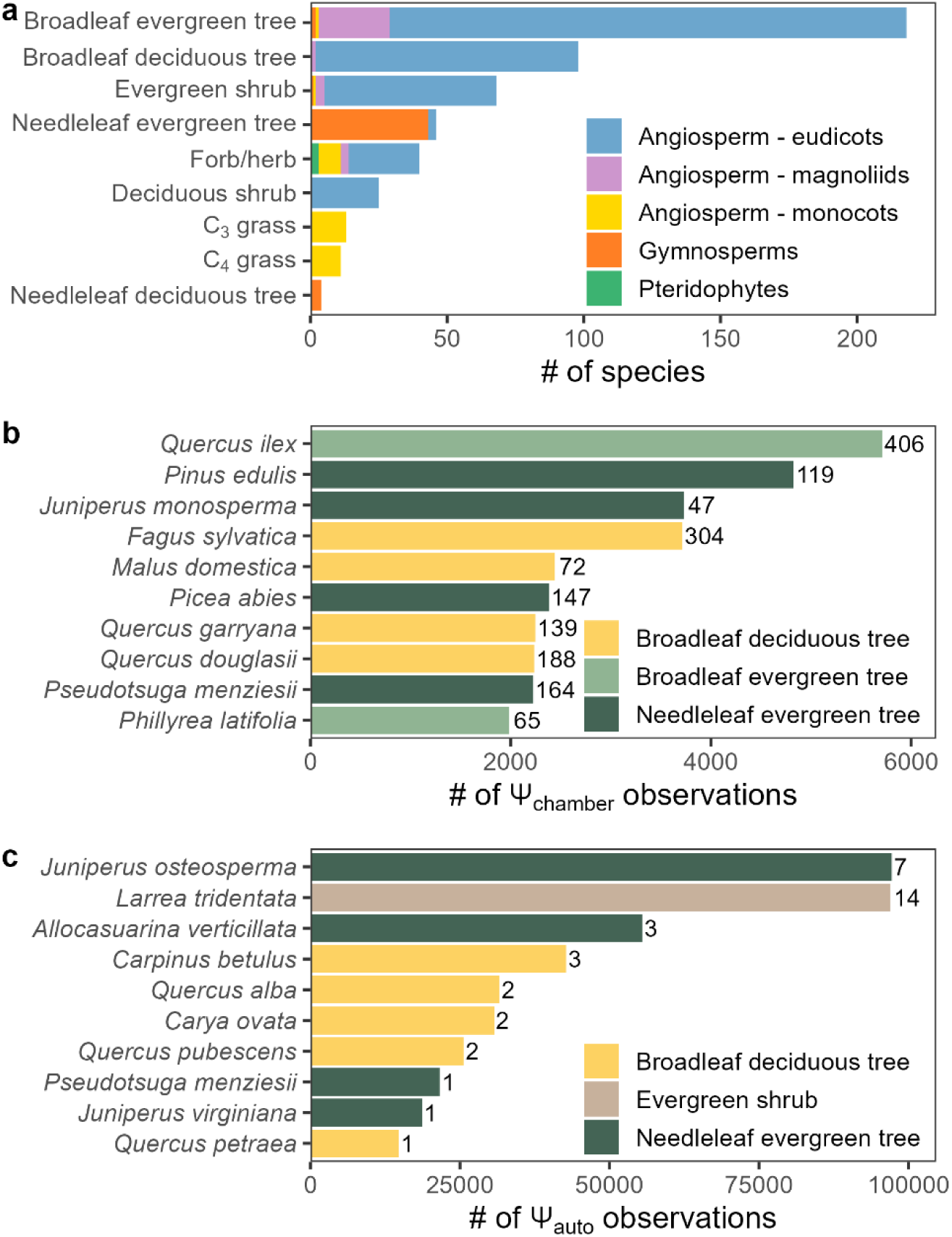
(a) Distribution of the species in the PSInet database by PFT and phylogenetic group. Top 10 species represented in the database by number of observations for (b) chamber, Ψ_cℎamber_, and (c) automated data, Ψ_auto_. Bars are labeled with the total number of individual plants pooled for each species. Note that while Ψ_auto_ has a much larger number of observations (two orders of magnitude), Ψ_cℎamber_ measurements were conducted on a larger number of individuals (one to two orders of magnitude).

Soil moisture and meteorological variables were encouraged but optional for all PSInet submissions. Data providers had the option to report soil water content or soil water potential at the individual, plot, or site level, and to submit measurements at two depths, “shallow” or “deep,” with the exact depth recorded as metadata. Most “shallow” soil moisture measurements were made between 10-25 cm, while the median depth of “deep” soil moisture was >60 cm. Shallow soil water content at the site-level was by far the most frequently submitted ancillary soil data (Fig. S3a). At least ten of the sites are part of the Regional Environmental Soil Hydraulics (ReESH) network (Crookshanks et al. *under review*), which openly shares soil hydraulic data including water retention curves and can be used to infer soil water potential from observed soil moisture content time series. An additional 46 sites indicated the existence of locally developed water retention curves that are not part of ReESH.

Meteorological variables, including temperature (T_air_), precipitation (Precip), relative humidity (RH), vapor pressure deficit (VPD), photosynthetically active radiation (PAR), and wind speed (Wind) were optional variables and were primarily measured locally within or near the site (e.g. above the canopy, in a clearing <1 km away); for a small number of datasets (n = 8), one or more meteorological variables were reported from off-site (> 1 km away). Air temperature was the most commonly reported variable (Fig S3b), available for ∼57% of the PSInet datasets (n = 164). Approximately 56% of the data submissions included RH and/or VPD (n = 160), and ∼30% included data on PAR (n = 86). Precipitation was reported for ∼46% of the datasets, and wind speed for ∼36%. Overall, 78% of the PSInet datasets located in flux tower or SAPFLUXNET sites reported at least some of the meteorological variables. Additional meteorological data from these sites may be available from FLUXNET (https://fluxnet.org/fluxnet-data-system/) and associated regional tower networks (e.g. AmeriFlux: https://ameriflux.lbl.gov/; OzFlux: https://www.ozflux.org.au/, Mexflux: https://mexflux.gitlab.io/) and SAPFLUXNET (https://sapfluxnet.creaf.cat/), though users are advised to carefully consult the data-use policies of these other networks. The percentage of PSInet sites that were not part of flux tower or SAPFLUXNET networks but nonetheless submitted meteorological data is lower than the percentage of sites that were part of these networks (∼49%). For a summary of additional data available for each dataset, see Table S3.

The PSInet database contains data collected between 1991 and 2025, though the majority of observations were made in the 2020s (Fig. 4a). There was a tradeoff between study duration and Ψ measurement frequency for the chamber data (Fig. 4b), which comports with the labor-intensity of chamber measurements. However, this pattern does not necessarily hold for automated Ψ data. Although maintaining instruments on living plant tissues is not trivial (see Restrepo-Acevedo et al. 2026), automated instruments hold promise for increasing data density without much compromise in study duration.

**Fig. 4.**
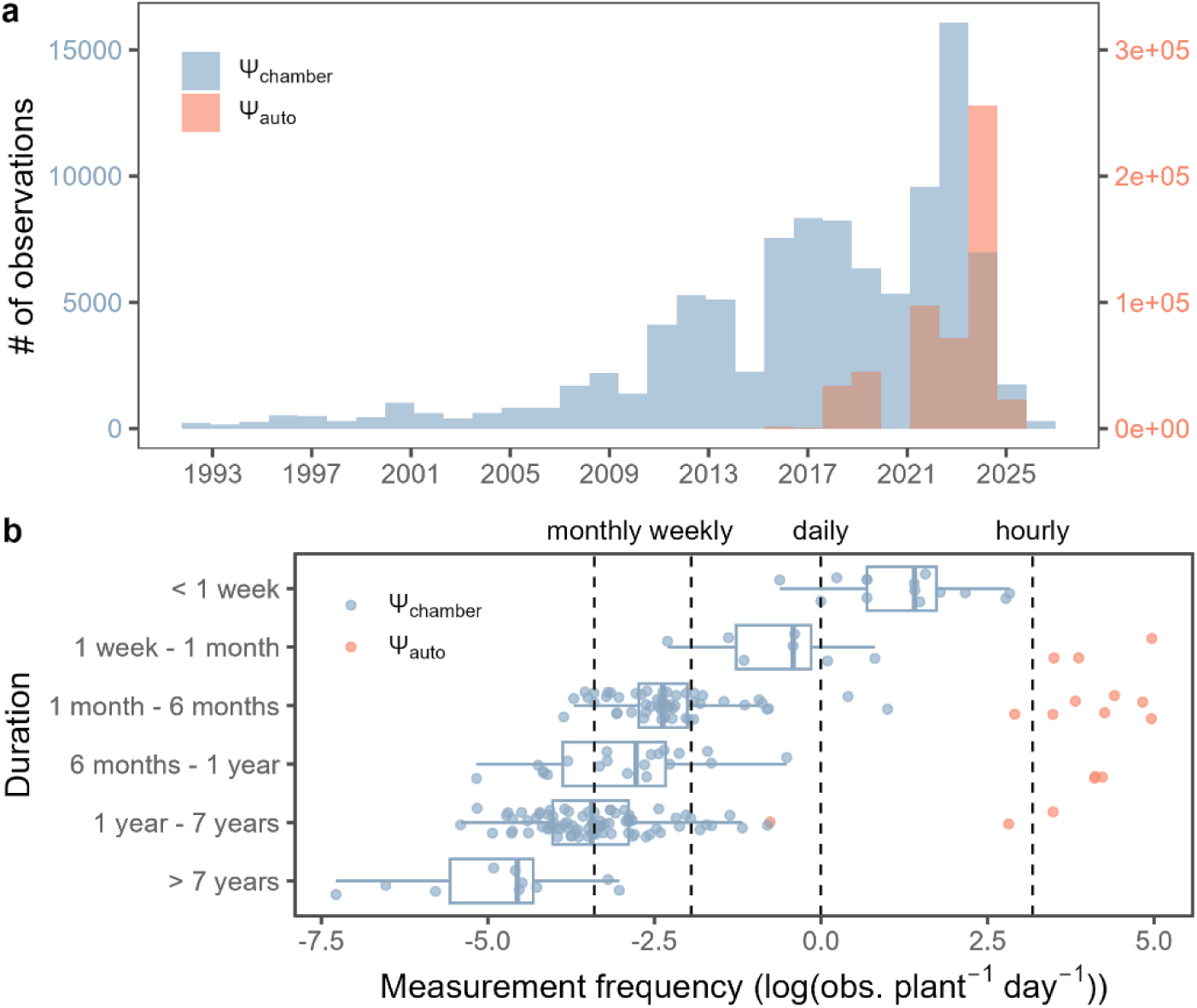
(a) Number of observations over time for chamber (Ψ_cℎamber_) and automated (Ψ_auto_) methods. (b) Relationship between study duration and mean frequency of Ψ measurements across bins of study duration. Note the log scale of the x-axis; values greater than zero indicate more than one measurement was taken per plant (or experimental unit) per day. Boxplots summarize Ψ_cℎamber_ data only.

### 4.1 Early insights from the PSInet database

In this section, we present preliminary results that demonstrate some of the research opportunities that can be enabled by the PSInet database. Many of these applications were originally proposed in Restrepo-Acevedo et al. (2024) and/or emerged during expert discussions at a workshop held in June 2025 at Indiana University – Bloomington.

### 4.1a Advancing conceptual frameworks for plant water use “strategies”

Classifying plant responses to drought and heat stress into generalizable ‘water-use strategies’ has been a hallmark eco-physiological research goal for decades. Initial categorizations were broad and frequently binary (e.g. ‘conservative versus profligate,’ ‘tolerant versus avoidant’), though recent schemes are more quantitative and continuous categorizations that link measurable traits to water flows and potential (Kannenberg et al 2022; Martinez-Vilalta et al. 2017; McCulloh et al. 2019; Meinzer et al. 2016). For example, one of the most popular modern frameworks is the concept of isohydry, which describes the degree to which plants regulate Ψ. More ‘isohydric’ plants close stomates quickly during drought which minimizes reductions in Ψ (but at the expense of reduced photosynthesis and a loss of transpiration cooling). More anisohydric plants keep stomates open longer, enabling more gas exchange but resulting in more rapid declines in Ψ that may increase the risk of drought-driven mortality (see additional discussion in Section 4.1d).

The slope of the relationship between pre-dawn and midday leaf water potential (*σ*) is one metric by which isohydry is commonly estimated (Konings et al. 2017; Martinez-Vilalta et al. 2014), though others exist (Klein et al. 2014; Meinzer et al. 2016; Salvi et al. 2022). The approaches relies on the common assumption that *Ψ* equilibrates throughout the soil-plant-atmosphere continuum during pre-dawn hours, such that pre-dawn Ψ serves as a proxy for soil water potential, which is otherwise difficult to measure (Crookshanks et al. *under review*). While the *σ* is frequently interpreted as a plant “trait,” in reality, it varies considerably among populations of the same species growing in different locations, and even within time for a single plant (Feng et al. 2019; Guo et al. 2020, Hochberg et al. 2018). This likely reflects a range of confounding factors including changes in plant hydraulic conductance (Martinez-Vilalta et al. 2017), declines in mid-day leaf *Ψ* driven by elevated VPD (Bourbia et al. 2025, Mencuccini et al. 2024, Novick et al. 2019), and increases in stomatal conductance during periods of excessive heat stress to increase evaporative cooling (Sicangco et al. 2026). However, in the absence of a representative, accessible database of Ψ time series, it has been difficult to integrate these factors into a more holistic framework (Kannenberg et al. 2022; Novick et al. 2022) and the concept of isohydry remains somewhat controversial (Hochberg et al 2018; Martinez-Vilalta et al. 2017).

Because the PSInet database directly addresses this data gap, it has the potential to advance the next generation of water use strategy frameworks. A natural first step is an exploration of the extent to which *σ* is conserved within species growing in diverse habitats. The PSInet database is well suited for this task, because many species in the database are represented by Ψ time series from more than one site. For three of the most well-represented species in the database (*Fagus sylvatica, Pinus sylvestris, and Quercus ilex*), we ran two sets of regressions: 1) a multiple regression of Ψ_MD_as a function of Ψ_PD_, which produces slopes (σ_i_) for each dataset, and 2) a simple linear regression between Ψ_MD_and Ψ_PD_regardless of dataset, which produced a combined σ. The *σ_i_* indeed varies between datasets within a single species, although to different degrees (Fig. 5a-c). For *F. sylvatica*, σ_i_varied marginally between datasets (p = 0.086), while for the *P. sylvestris* and *Q. ilex*, σ_i_varied significantly between datasets (p < 0.001). The combined σ explains 68% of the variation for the relatively anisohydric *Q. ilex* (combined *σ* = 0.65) and 40% for *P. sylvestris* (combined *σ* = 0.74). *F. sylvatica,* in contrast, is the most isohydric of the three species (combined *σ* = 0.42), and as expected for isohydric plants, the cross-site relationship between predawn and midday water potential is weaker (R^2^ = 0.19). However, we note that iso-/anisohydric classification should be interpreted with some caution, as population-specific drought responses may also be strongly shaped by other pedo-climatic factors that can vary from one site to the next.

**Figure 5:**
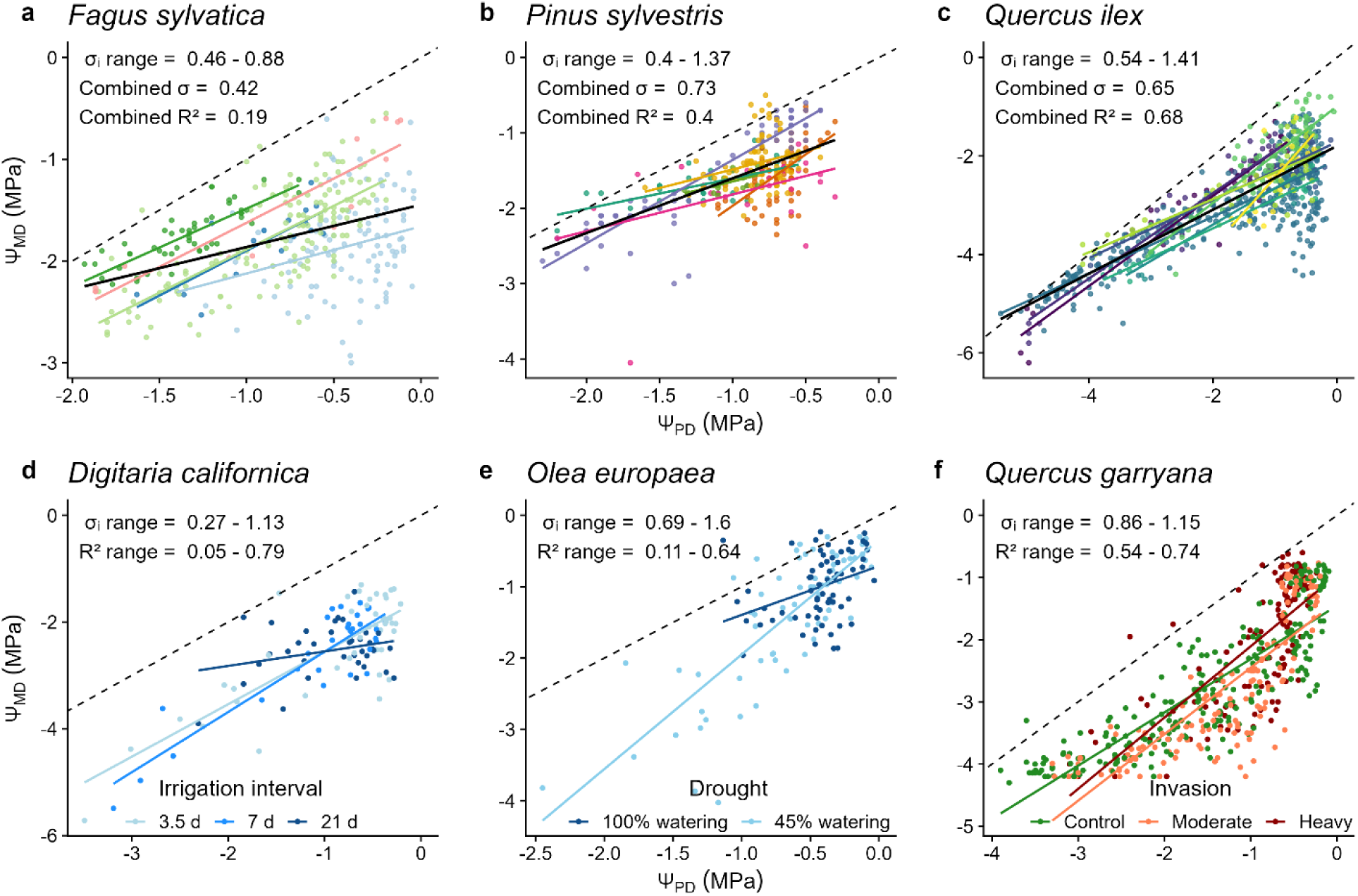
(Top row, a-c): The relationship between midday and predawn Ψ for the three of the top 10 most prevalent species in the PSInet database, with each color representing a different dataset. Colored lines show the within-dataset lines of best fit determined from linear regression, with σ_i_ representing the dataset-level slope. The solid black line shows the cross-site line of best fit for the whole species, with an associated ‘combined σ’ across datasets and R^2^. (Bottom row, d-f): The relationship between midday and predawn Ψ for the three species subjected to different treatments, with each color representing a different treatment. (d) Digitaria californica was irrigated at three different intervals in Guo_2, (e) Olea europaea was subjected to drought in ESP_SAN, and (f) Quercus garryana was measured in plots with no, moderate, or heavy invasion by Pseudotsuga menziesii in Ker_3. Colored lines show the within-treatment lines of best fit and σ_i_ represents the treatment-level slope. The range of R^2^ across treatments is also displayed. The dashed black line represents the 1:1 line.

A similar multiple regression was performed on three datasets describing plants of the same species that experienced different experimental treatments. In this analysis, σ_i_represents the slope of each treatment within the dataset (Fig 5d-f), but a combined σ was not calculated. While species-level differences in *σ* were evident (e.g. the σ range of *Quercus garryana* and *Olea europaea* are narrower than that of the C4 grass *Digitaria californica*), so too were differences in the *σ* driven by treatments that altered water availability (*D. californica, p < 0.001* and *O. europaea, p = 0.002*) and community composition (*Q. garrryana*, p < 0.001).

Additional information contained in the PSInet database and other environmental networks has the potential to allow us to better understand the mechanisms underlying these dynamics. A useful starting point is a simple steady-state representation of the linkages between soil Ψ (Ψ_s_), the leaf Ψ (Ψ), whole plant transpiration (*T*_r_), canopy-level stomatal conductance (*G*_s_), and whole plant hydraulic conductance (*k*) that emerges from Darcy’s Law (Whitehead et al. 1981; McDowell et al. 2015):

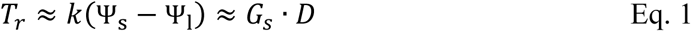

where *D* is the vapor pressure difference between the leaves and the surrounding air, which is frequently assumed to be equal to the vapor pressure deficit of the air (e.g. the VPD, but see Cernusak et al. 2024; Novick et al. 2024). We also retain the imperfect assumption that predawn measurements of Ψ_1_represent the root-zone soil water potential (see additional discussion in Section 4.1c).

From Eq. [1], the slope of the relationship between *T*_r_and the difference between pre-dawn and midday Ψ (hereafter ΔΨ) provides a dynamic estimate of the plant-scale *k*. Many site-level studies have used this approach to understand how environmental conditions and ontogeny drive variations in *k* over time (see, for example, Zhao et al. *under review*), using both tree-level sap flow data and, more recently, stand-level estimates of transpiration derived from flux towers (Wood et al. 2023).

Because 110 PSInet sites are also nodes of SAPFLUXNET or FLUXNET, a more systematic evaluation of the spatio-temporal dynamics of *k* during drought and heat stress is possible. As an illustrative example, here, we used sap flow and leaf Ψ data for all sites that are represented in both SAPFLUXNET and PSINET. We calculated tree-level sap flow per basal area at midday for the SAPFLUXNET datasets by averaging hourly or sub-hourly sap flow per basal area between 11:00 and 14:00 solar time. We then calculated a 95% quantile of this midday value, as an estimate of maximum sap flow normalized by tree size which was then averaged at the site and species level. For the same sites and species, we calculated ΔΨ and estimated the maximum ΔΨ. The relationship between the two variables was, as expected, positive. However, substantial scatter suggests the important influence of spatial and temporal dynamics of *k*, within and across species (Fig. 6). A more granular analysis could track the relative variation in plant-level *k* for individuals or stands over time—information that could then be cross-referenced against the expected variations in *k* obtained from xylem vulnerability curves and information in the XFT database. Other opportunities include an expansion to the ecosystem scale by exploring coordination between measured tower-derived transpiration estimates and the ΔΨ (e.g. Wood et al. 2023). Additional discussion on approaches for scaling from the plant level to the ecosystem can be found in section 4.e.

**Fig 6:**
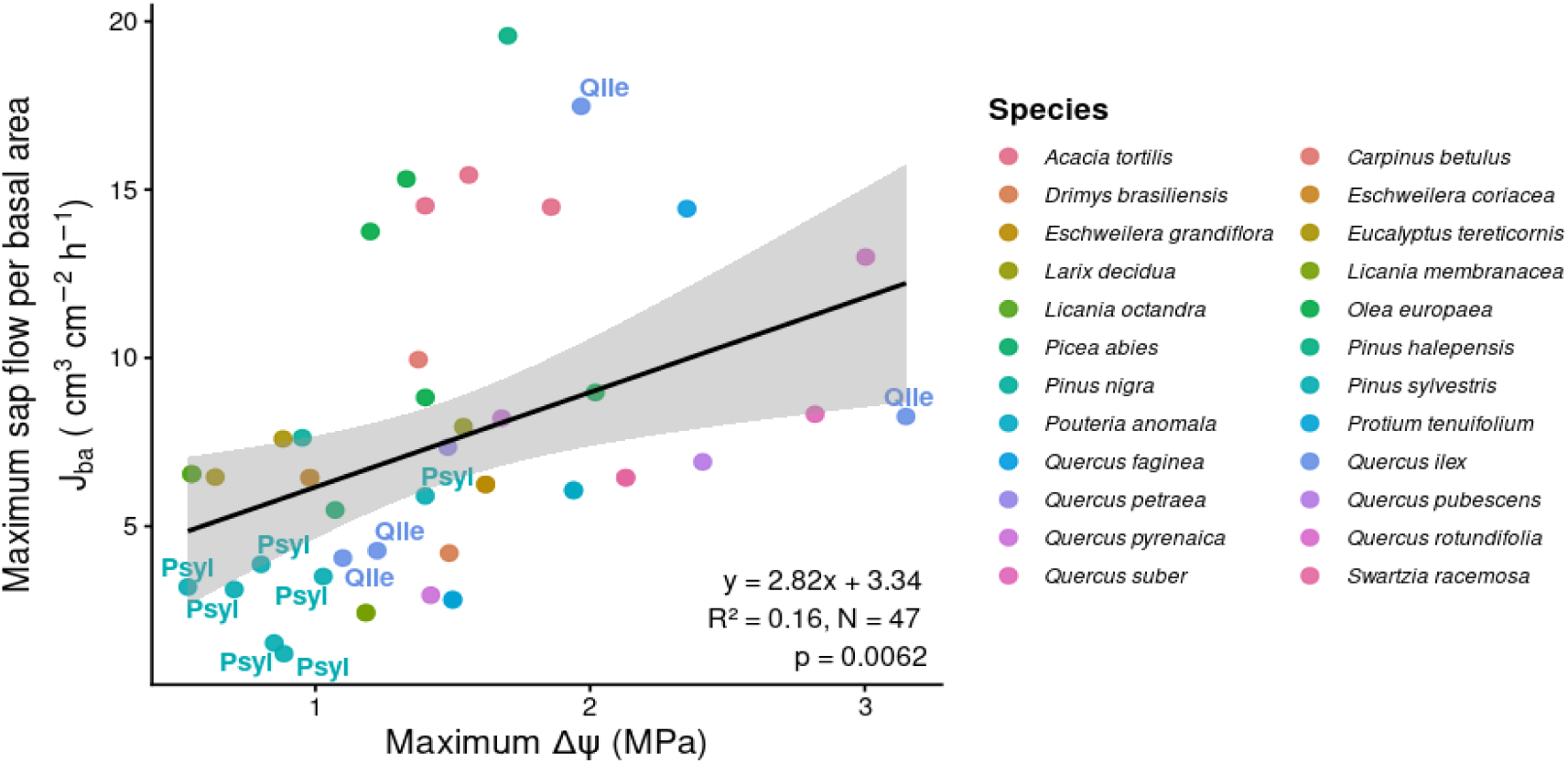
The relationship between maximum sap flux rates and maximum ΔΨ across 47 combinations of site and species tree species. Data for Pinus sylvestris and Quercus ilex, two of the most well-represented species in SAPFLUXNET and PSInet have been labelled in the plot.

### 4.1b Disentangling the influence of co-evolving environmental drivers on plant *Ψ*

Plant responses to drought stress have historically been linked primarily to changes in soil water availability, despite the fact that elevated VPD during drought can independently suppress midday *Ψ* (Bourbia et al. 2025, Mencuccini et al. 2024; Novick et al. 2019). This is evident by transforming the right side of Eq (1) into: 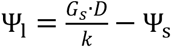. Climate change is driving large increases in VPD and air temperature almost everywhere (Novick et al. 2024), but the impacts on precipitation and soil water are more spatially heterogeneous (Cook et al. 2014). As a result, the relationship between soil and atmospheric drought is changing, and understanding how plants respond to each is a key research goal (Grossiord et al. 2020). Prior work has focused on disentangling the impacts of soil water versus VPD on ecosystem fluxes measured by sap flux and flux towers, and on proxies for plant function observed from remote sensing (Flo et al. 2023; Fu et al. 2025; Lowman et al. 2023; Novick et al. 2016). However, until now, the lack of a centralized repository of plant Ψ timeseries has made it difficult to empirically quantify the impacts of VPD on Ψ.

Because soil water and VPD tend to be relatively decoupled at hourly and daily timescales (Novick et al. 2016), disentangling their impacts on Ψ is easiest when using data collected at a high temporal resolution (e.g. hourly or daily) and that span meaningful gradients in both drivers. In the PSInet database, approximately 1/3 of the chamber time series report plant Ψ recorded repeatedly (at least 4 times) over the course of at least 5 days, which could be sufficient to isolate VPD impacts. However, the long-term continuous water potential data are especially well-suited for this task. Preliminary analysis of two automated datasets in PSInet shows that, after controlling for soil water, a clear inverse relationship between Ψ and VPD emerges (Fig. 7), especially when soils are dry. Future work could synthesize continuous Ψ data to understand, for example, if plants that are more an/isohydric with respect to soil water are also more an/isohydric with respect to rising VPD. Other work could focus on novel approaches for disentangling the impacts of rising VPD from those of excessive temperature, since the latter can impact plant hydraulic functions in ways that are unique from the impacts of soil water and VPD (Aparacido et al. 2020; Drake et al. 2018; Ruehr et al. 2019; Still et al. 2023).

**Fig. 7:**
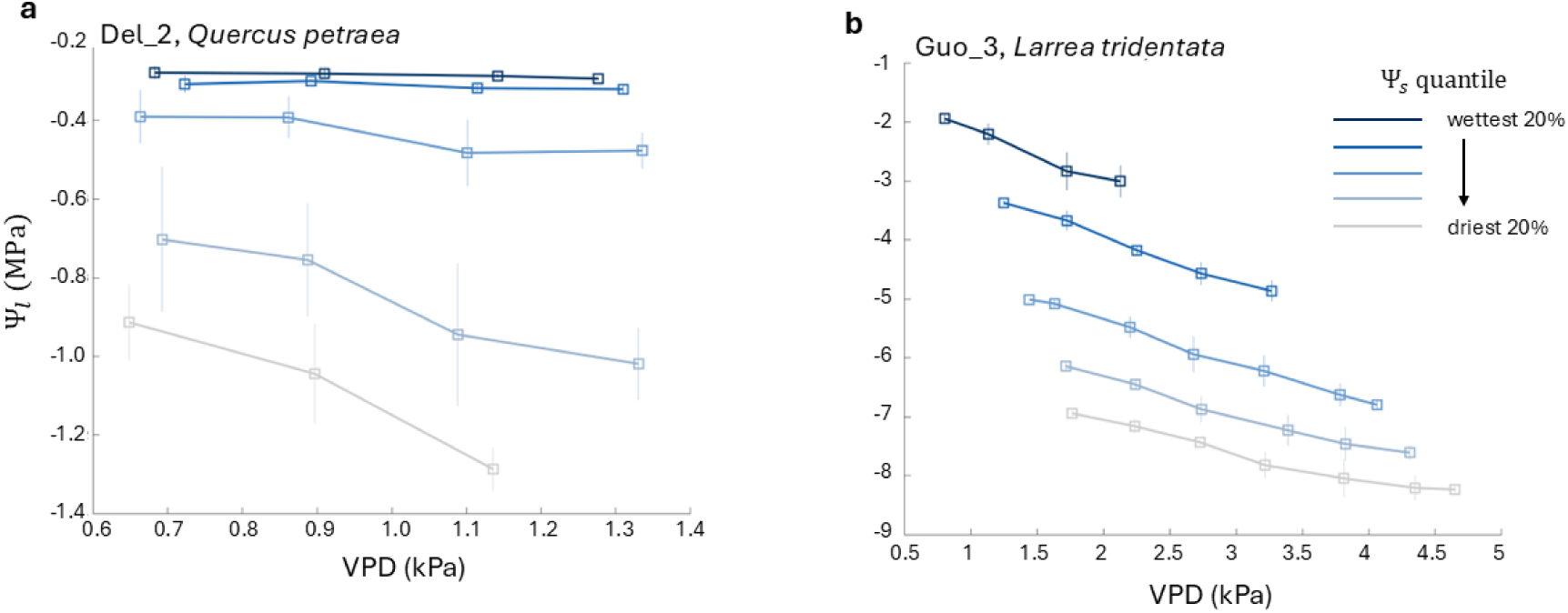
The negative relationship between stem Ψ and VPD for two PSInet sites (Del_2 and Guo_3) that contributed automated data. The data are binned into five quantiles on the basis of soil water availability (using pre-dawn water potential as a proxy for soil water potential). In an arid site dominated by Larrea trudebtata (Guo_3, panel b), the sensitivity of Ψ to VPD is evident across all soil water bins, but tends to diminish when soil is especially dry. In contrast, in the more mesic site (panel b), the sensitivity of Ψ to VPD increases as soil dries.

### 4.1c Revisiting the long-held assumption of nocturnal equilibration

The widely used assumption that plant and soil Ψ equilibrate in predawn hours is largely an assumption of convenience, as it is challenging to measure root-zone soil Ψ through other means. In reality, these assumptions will not always hold. For example, eco-physiological processes like nocturnal transpiration, capacitance, changes in hydraulic resistance, and hydraulic insolation from the soil can all prevent predawn equilibrium (Buccci et al. 2005; Donovan et al. 2002) and cause *Ψ*_L_to be lower than the root-zone soil water potential. This disconnect, in turn, complicates efforts to understand the relationship between plant response and soil water status. The continuous *Ψ* data aggregated by PSInet offer a unique opportunity to better understand how often this assumption holds (or fails). Specifically, if pre-dawn equilibrium does occur, then we would expect the measured *Ψ* to be relatively constant in the hours leading up to dawn. However, when conditions fail to equilibrate, then *Ψ* could continue to increase throughout the pre-dawn hours without ever achieving a stationary value. Preliminary analyses from three species in the PSInet database suggest that non-stationary *Ψ* in the hours preceding dawn is in fact quite common (Fig. 8), and should motivate additional analyses to understand the conditions that promote disequilibrium, and the consequences for our understanding of soil water potential.

**Fig. 8:**
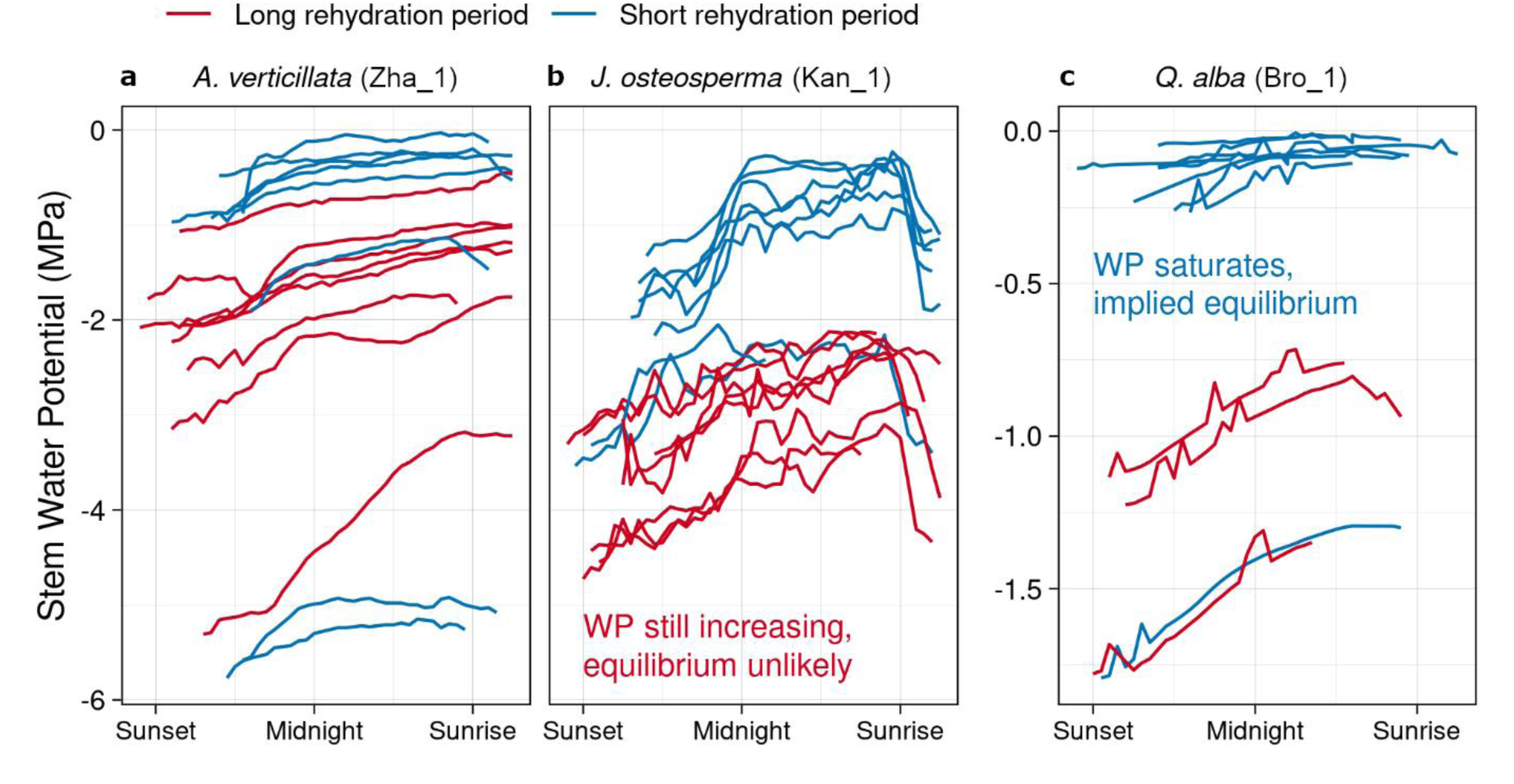
Stem Ψ measured every half-hour for three species (Allocasuarina verticillata, Juniperus osteosperma, and Quercus alba) at three sites, shown on a relative time scale. Each line is a different nocturnal period, and together they reveal how the shape of the overnight rehydration curve is related to pre-dawn equilibrium (or lack thereof) between plant and soil water potentials. A simple model of dynamic stem water recharge can be fit to the continuous Ψ_stem_ measurements to estimate the length of time required for rehydration. When this length of time is longer than the duration of nighttime (red curves), the plant may not have fully equilibrated with the soil water potential. The PSInet site code is indicated above the panels after the species name.

### 4.1d Understanding the risk of drought-driven hydraulic failure

Hydraulic failure – or the cessation of xylem water transport due to embolism – is broadly recognized as one of the most important mechanisms that can trigger drought-induced mortality in plants (Choat et al. 2018; Hammond et al. 2020; McDowell et al. 2008). The risk of hydraulic failure is often quantified by the hydraulic safety margin, which represents the difference between the minimum plant Ψ (Ψ_min_) and the plant Ψ associated with a 50% loss in hydraulic conductivity (i.e., the P_50_Meinzer et al. 2009). The latter is derived from vulnerability curve measurements made in the laboratory, and the P_50_of many plant species is available from XFT. While XFT also aggregates user-provided information on the minimum Ψ, these values are highly sensitive to the environmental conditions that occurred during the course of any given study. For example, a study that occurred in the absence of drought will likely report a much higher Ψ_min_ than a study that took place during an extreme drought or heat event. Data from PSInet illustrate the problem of assigning a species-level Ψ_min_ trait in this way, as the reported Ψ_min_ among time series of the same species is quite large (Fig. 9).

**Fig 9:**
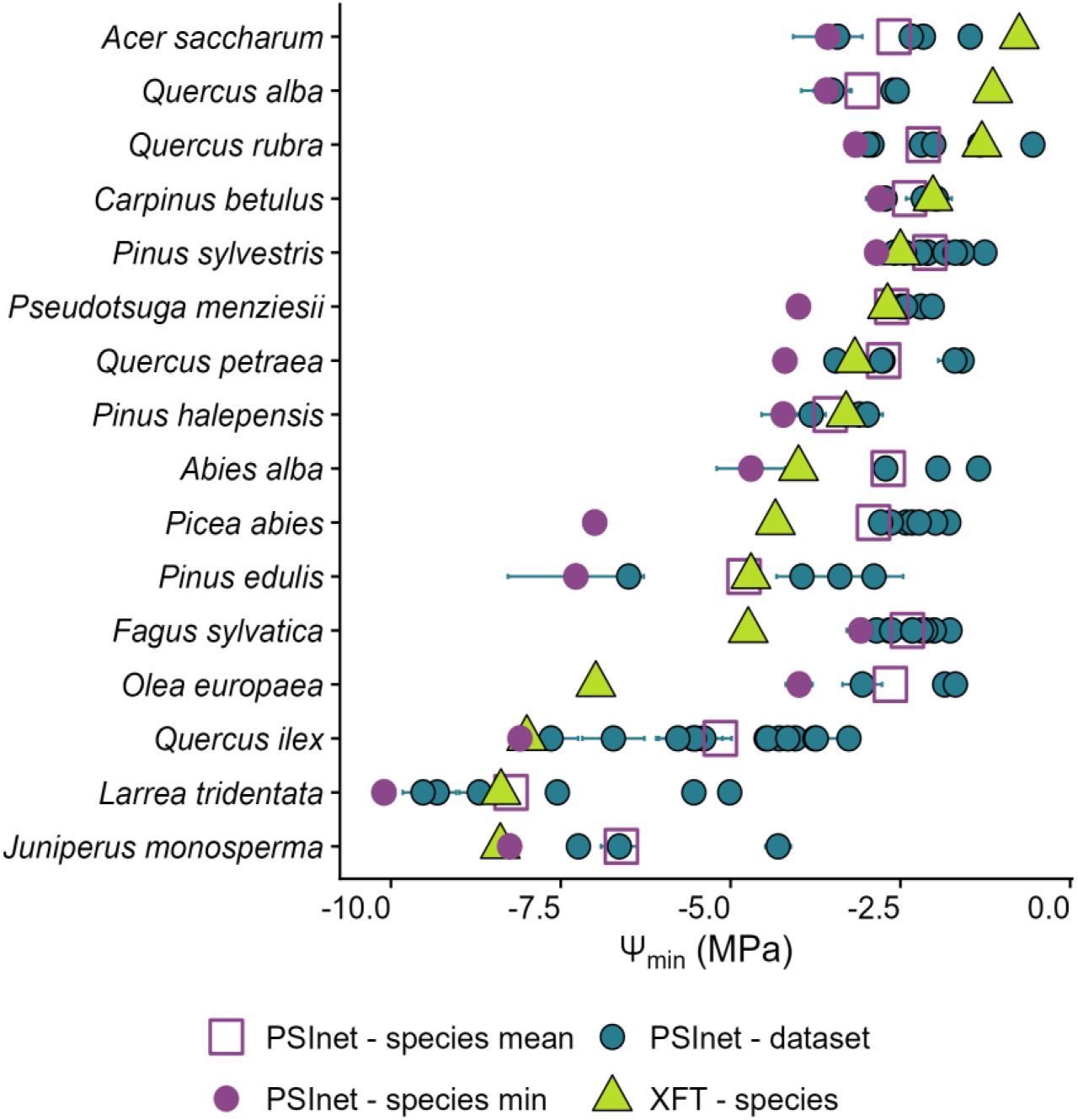
The minimum water potential for a subset of species within the PSInet database, using chamber data only. Each circle represents the mean of the lowest four values in one dataset; the error bars represent the standard deviation around the mean; only datasets with 30 or more measurements were included. Purple circles represent the lowest minimum water potential across datasets based on PSInet data; open squares represent the mean minimum water potential across datasets based on PSInet data; green triangles represent the lowest minimum water potential as reported in XFT.

Moreover, because Ψ_min_ represents the tail end of a distribution of plant Ψ, its determination is sensitive to sample size effects even if the study period represents a range of hydrologic conditions. Recent work by Martínez-Vilalta et al. (2021) highlights the usefulness of extreme value theory for circumventing these challenges. Extreme value theory is often used by engineers, for example, to estimate the magnitude of a 50- or 100-year flood event given a time series that may not necessarily capture that event. The same principle can be applied to plant water potential time series to estimate the lowest value of Ψ_min_ that a plant might experience during a given period of time. However, a systematic evaluation of Ψ_min_ derived from extreme value theory first requires a database of representative water potential time series. With the release of the PSInet database, the door is open to more rigorous evaluation of Ψ_min_ which may have the potential to reshape our understanding of which species are most vulnerable to drought-driven hydraulic failure.

### 4.1e Integration with remote sensing products and land-surface models

The previous sections highlight knowledge gaps that can largely be addressed through synthesis of plant-level data, for which the PSInet database is particularly well suited. However, many of the most pressing questions in plant drought research require an ecosystem- or landscape-scale perspective. Robust benchmarking of satellite-derived proxies for plant water status requires ground truth data collected at scales (typically km^2^ or larger) that match remote sensing pixels (e.g., Tian et al. 2016). Likewise, the land surface models used to predict drought responses in space and time primarily operate at the ecosystem scale and need to be tested and validated with ecosystem scale data (e.g., Corak et al. 2024; Lowman & Barros 2018). This is also the scale at which flux towers operate, and fully harnessing the potential for integration across the dozens of PSInet datasets collected in the footprint of flux towers requires strategies to transform plant-level measurements into ecosystem scale time series.

Upscaling from plant to ecosystem scales is challenging. In biodiverse ecosystems, it can be difficult to representatively sample all species and canopy layers present. Even when the number of species is small, it is also difficult to develop sampling schemes that robustly capture soil and microclimate gradients. And in all ecosystems, processes like soil evaporation and canopy interception that act beyond the plant scale can conceptually decouple ET from plant-level measurements of water status and flows. A preliminary review of the PSInet data reveals that variability from one species to the next within a site can be substantial relative to variations in the site-to-site means (Fig. 10). As a result, attempts to upscale plant-level Ψ data to the ecosystem level must consider multiple species and be carefully executed in ways that incorporate uncertainty and the relative abundance of both measured and unmeasured species in a given site.

**Figure 10:**
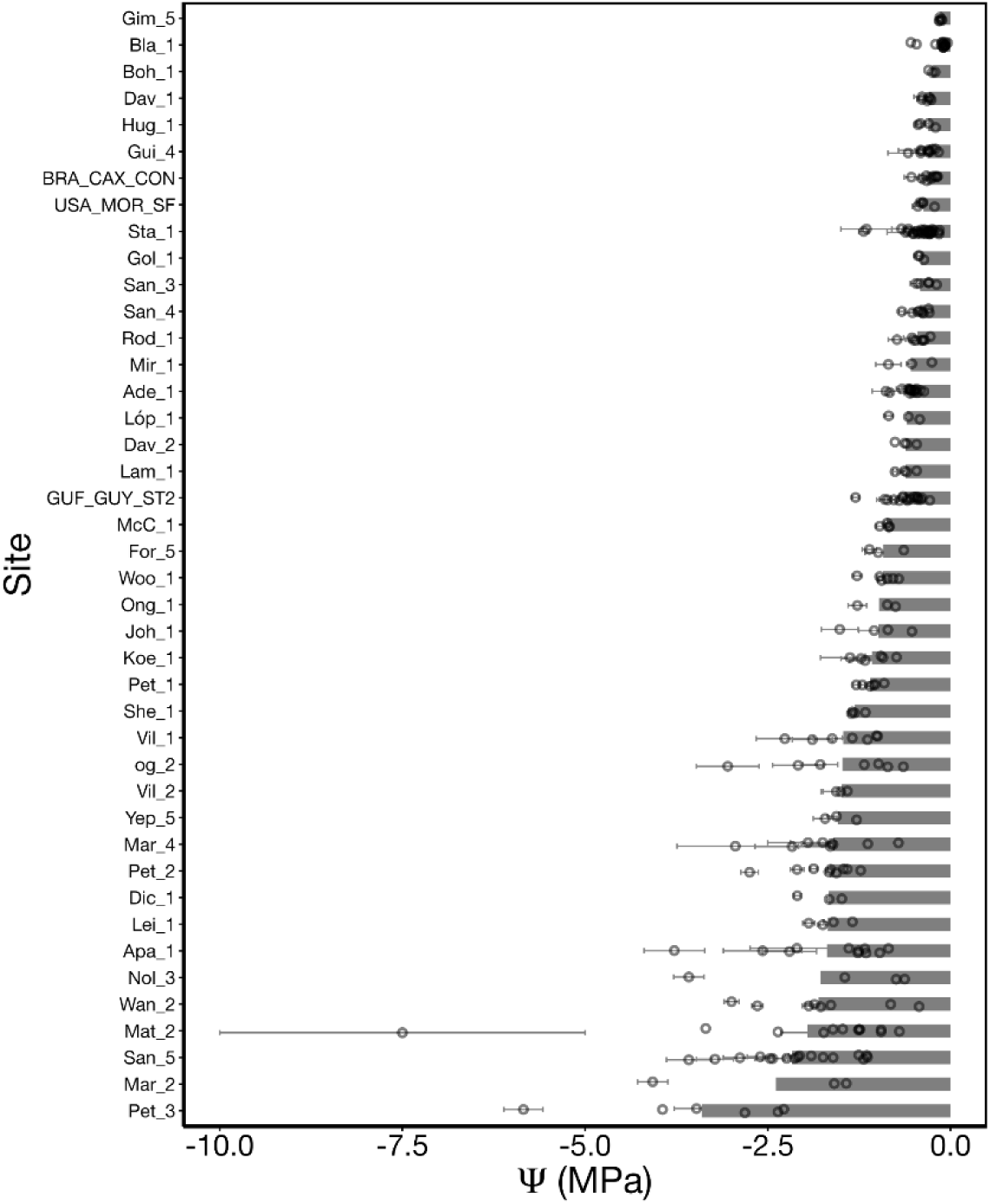
Site and species average predawn Ψ ordered by mean site water potential, for PSInet sites with greater than 3 species per site. Gray bars represent site-wide mean water potential and points represent the averages by species, with the bars representing standard error around the mean.

For applications involving flux tower data or remote-sensing products with a relatively small footprint, weighted averages of PSInet Ψ timeseries may be a feasible strategy to produce upscaled estimates of ecosystem water potential. For example, a preliminary analysis of PSInet data suggests that for many species, a direct relationship between ECOSTRESS ET (which is based on land surface temperature measurements at a 70-m resolution, Fisher et al. 2020) and ΔΨ is evident (Fig. 11). However, in other sites, the relationship is less consistent, which could indicate the confounding effects of scale-emergent processes or dynamic variations in hydraulic conductance. We note that satellite-based ET is also uncertain because satellites can only sense variables related to ET, and not ET itself, requiring semi-empirical modeling in the ET estimation (Pan et al. 2020). Future work could leverage a more representative set of PSInet time series to identify the conditions under which remotely sensed ET is most strongly coupled to plant water status.

**Fig. 11.**
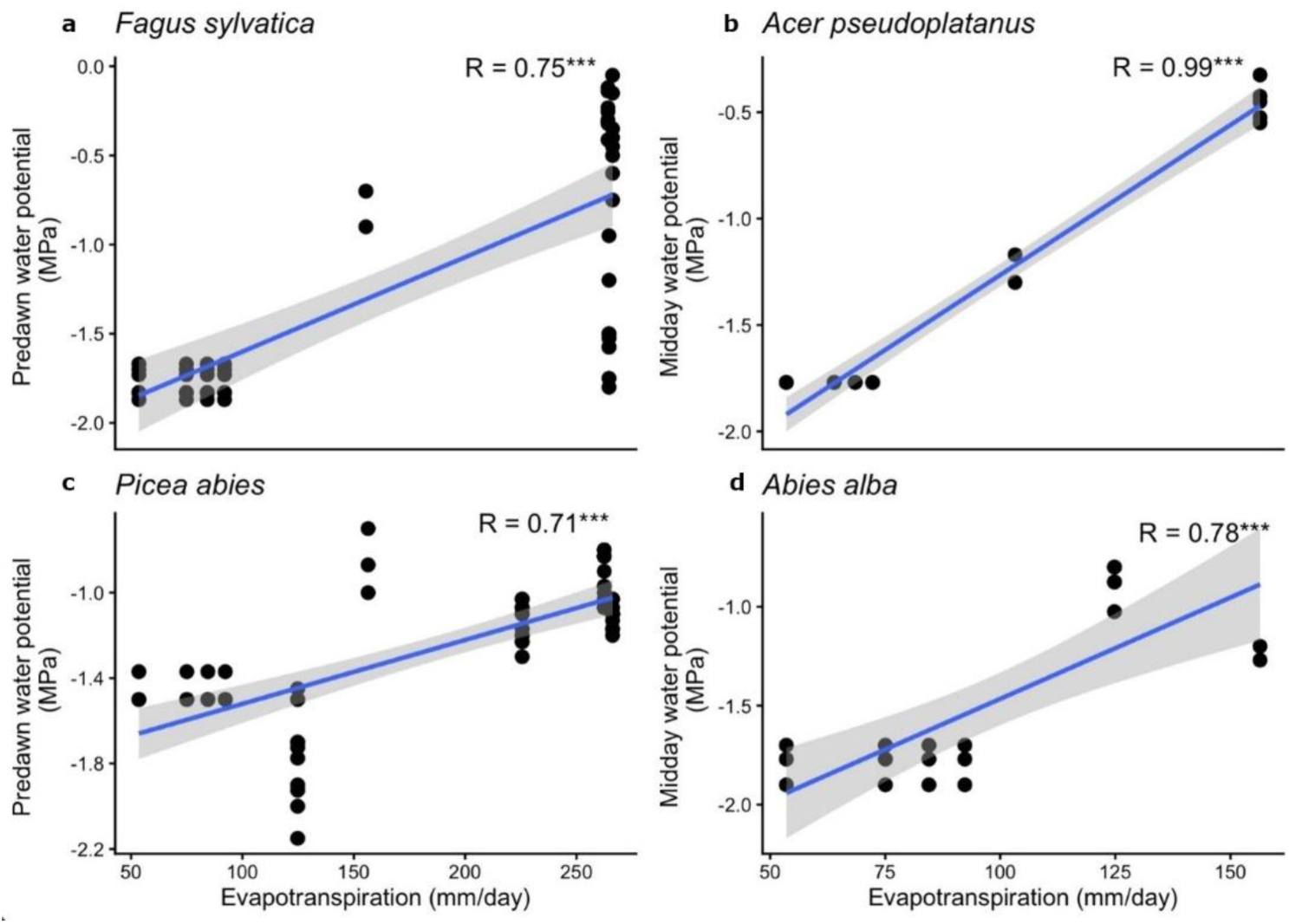
Pre-dawn (left column) and mid-day (right column) leaf water potential data from PSInet, plotted as a function of ECOSTRESS ET data for the same species at different sites and times.

Other exciting research questions require the integration of remotely-sensed datasets with a larger resolution, including satellite derived estimates of gross primary productivity, solar-induced fluorescence, and canopy water content derived from vegetation optical depth, or VOD (; Chapparo et al. *in press*; Konings et al. 2021; Magney et al. 2019; Turner et al. 2006). Comparing VOD with Ψ is of particular interest since previous research suggests VOD is potentially sensitive to plant water potential (Holtzman et al 2020; Konings et al 2021b). Further research is needed to enable retrieval of Ψ from VOD, which will likely be supported by the recent development of transmissometry GNSS-based systems for the in-situ estimation of vegetation optical depth (VOD) (Feldman 2024; Humphrey & Frankenberg, 2023).

For these coarser scale products, the utility of PSInet data will depend on the extent to which the measured species are representative of the larger area covered by the pixel footprint. This will vary by dataset and study, and may require additional engagement with the literature, communication with data providers, and/or information contained in regional- or national plant survey and inventory datasets. Nevertheless, data contained in PSInet may be useful for understanding how to design a spatially representative validation campaign. For example, it could be used to assess the amount of inter-specific and intra-specific variability in Ψ and how many replicates are needed to sample this variability. The PSInet database could also be used to study the degree to which extrapolations between species based on lineage are possible (Sanchez-Martinez et al. 2023, L. Anderegg et al. 2022).

Finally, the impacts of drought and heat stress remain a major unresolved source of uncertainty in land surface modeling (Mencuccini et al. 2019). More explicit representation of plant hydraulic processes, including plant water potential, has been proposed as a pathway for reducing this uncertainty (Anderegg et al. 2012), and plant hydraulics are increasingly being represented in land surface models (Christofferson et al. 2016; Kennedy et al. 2019; Mirfenderesgi et al. 2016; Xu et al. 2023). However, the scarcity of accessible observations of plant Ψ has made it difficult to benchmark and test these new schemes. Key unresolved issues include robust comparisons of different hydraulic model structures (Sabot et al. 2022) and critical evaluations of model parameterization schemes – for example, comparing model-data fusion approaches (Liu et al. 2021) to parameterizations grounded in theoretical principles (Sabot et al. 2022; Sperry et al. 2016). Addressing these challenges will benefit from more comprehensive *Ψ* datasets in PSInet, which could be strategically compared with or assimilated into models that explicitly represent the evolution of plant Ψ.

For example, although most land surface models represent an ecosystem using a single “average” plant, demographically enabled land surface models, such as ED2 (Medvigy et al. 2009) and FATES (Xu et al. 2023), explicitly track physiological dynamics across cohorts that share similar size and functional type. This cohort-based structure provides a more scale-consistent link to in situ measurements made on individual plants. Because different cohorts can respond to the same drought event in distinct ways (Fig. 12), this model structure creates new opportunities for scale-coherent model–data comparison and integration. PSInet presents unique opportunities where leaf water potential observations could be assimilated into, or used to evaluate, model outputs at their native observational scale, rather than being aggregated semi-arbitrarily to the ecosystem scale. In this way, cohort-resolved models may provide a more direct and scale-consistent constraint on plant hydraulic processes, improving our ability to diagnose drought responses mediated by diverse canopy composition and structure and, ultimately, to improve model representations of vegetation water stress.

**Figure 12:**
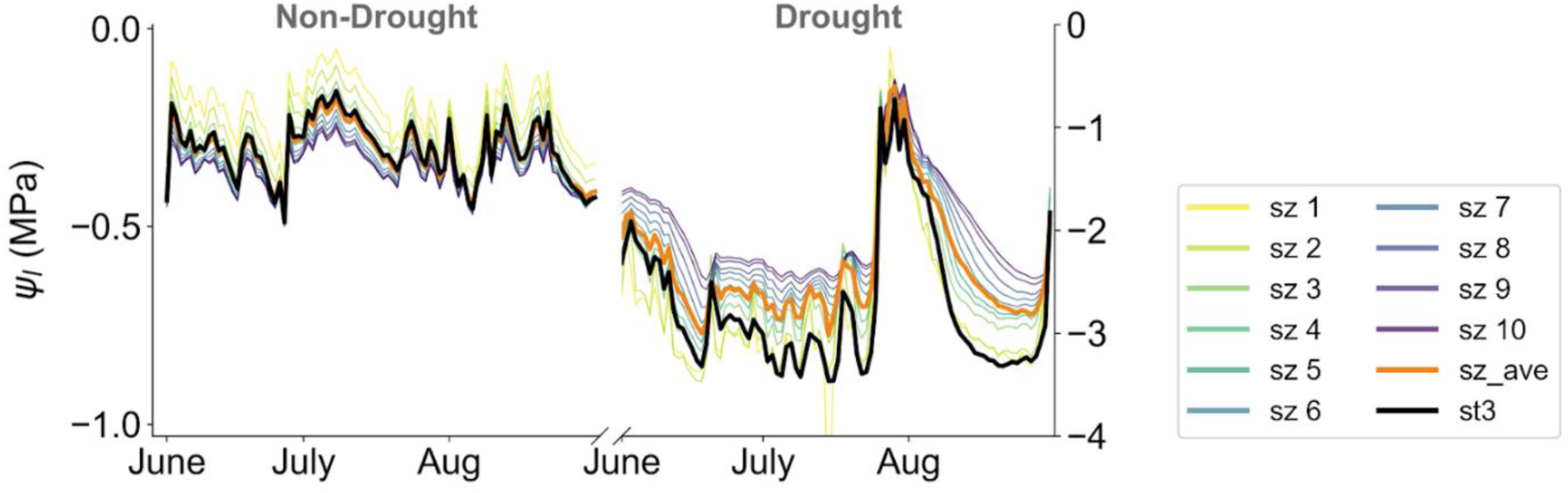
Time series of leaf Ψ produced by the ELM-Fates model for the US-Chr chestnut ridge AmeriFlux sites. Different lines represent different size classes. This figure demonstrates the capability of next-generation models that incorporate plant hydraulics to provide information about water potential dynamics that could be validated with PSInet data, alone or in combination with information from other environmental observation networks.

## Conclusion

Anticipating and preparing for life on a warming planet requires a predictive understanding of how increasing drought and heat stress will affect terrestrial plants and the many services they provide. The *Ψ* of soils and plants is a fundamental driver of ecosystem water flows, and directly controls many aspects of plant functioning during drought. However, observations of plant *Ψ* have historically been disjointed and inaccessible, constraining the synthetic research necessary to improve conceptual understanding and predictive models of plant drought responses. The PSInet plant *Ψ* database—which is the first global database of water potential time series—directly addresses this data gap. It includes data from 285 sites representing 523 species distributed across a broad range of biomes. Preliminary analyses demonstrate the potential of this database to better quantify key plant traits (e.g. *σ*, Ψ_min_) and their coordination, to isolate the impacts of declining soil water from rising vapor pressure deficit and temperature, and to critically evaluate the long-held assumption of pre-dawn equilibrium. We also show that integration of data from the PSInet database with other environmental networks can permit us to better understand and constrain hydraulic conductivity and evapotranspiration. Finally, we discuss strategies for upscaling plant-level PSInet measurements to the ecosystem scale where they can be used to benchmark and develop next-generation remote sensing data products and land-surface models. The PSInet database provides a foundation upon which the scientific community can build towards a more mechanistic understanding of plant responses to their environmental conditions. The work made possible by this dataset will better equip us to predict how vegetation will adapt to changing climate conditions across scales.

## Acknowledgements

The authors gratefully acknowledge the global community of scientists, including students and research technicians, who generated the data that form the foundation of the PSInet plant water potential database. We also thank the National Science Foundation— Division of Integrative Organismal Biology for funding the work of PSInet via a Research Coordination Grant (#2243900). We also acknowledge the Fluxnet Coordination Project (NSF Accelnet Grant #2113978) and Indiana University – Bloomington for providing additional funding to support a workshop held in Bloomington in June 2025, from which many ideas for this paper emerged. We acknowledge the author-level data contributions of Joel A. Biederman to Guo_2 and Russell L. Scott to Ham_6 and Ham_7; circumstances prevented their inclusion as full authors on this paper.

## Author contributions

J. Guo and K. Novick conceived of the study and the development of the PSInet network and related activities, in close collaboration with D. Johnson, K. McCulloh, and J. Nippert. J. Guo, A.M. Restropo-Acevedo, and M. Browne designed the digital infrastructure of the database. J. Guo, M. Browne, K. Novick A. Endsley, L. Sack, R. Poyatos, N. Vinod, and Y. Liu contributed figures and text to the manuscript, and together with S. Kannenberg, A. Konings, A. Feldman, J. Dukes, D. Johnson, L. Sack, W. Hammond, J. Green, L. Lowman, D. Beverly, K. Hultine, and J. Hie, provided substantial input on early versions of the text. Other co-authors, including E. Bucior, D. Chaparro, A. Crookshanks, X. Feng, R. Freitas, J. Green, B. Harrison-Day, K. Hultine, M. Kant, J. Missik, K. Morris, K. Ogle, A. Pivaroff, M. Rao, T. Robinett, K. Shellenberg, C. Valle Rodriguez, and Y. Zhang contributed to the scope of the paper, and in particular section 4, though their participation at a workshop held in June 2025 at Indiana University – Bloomington. All other authors shared data to the database and participated in multiple rounds of review of their data contributions. All co-authors reviewed the manuscript.

## Data Availability

The code that generated the PSInet database are available at https://github.com/PSInetRCN/data-qa and https://github.com/PSInetRCN/PSInetdb. Data can be accessed through the R package ‘PSInetR’ (https://github.com/PSInetRCN/PSInetr) as a database object or Zenodo as rectangular .csv files (10.5281/zenodo.21347356). All links will be made publicly accessible by October 1, 2027. Information on the data use policy can be found here: https://psinetrcn.github.io/data_terms.html.

