## Supplementary Data and Tables for "The PSInet Plant Water Potential Database: advancing new perspectives on plant water status, traits, and hydraulic processes"

**Supplementary Figures and Tables**

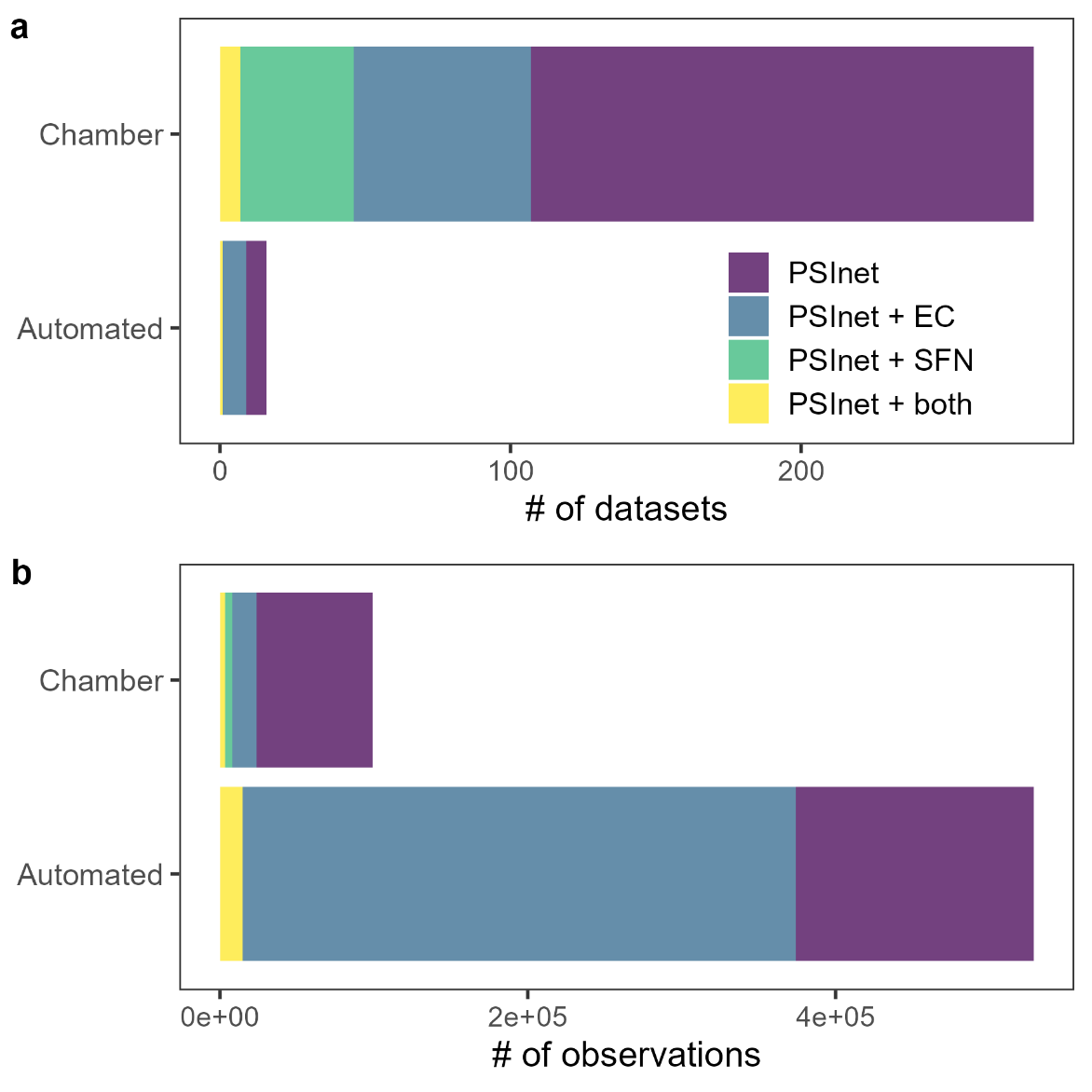

*Figure S1. Number of a) datasets and b) observations by chamber versus automated measurement types, colored by co-location with other data networks.*

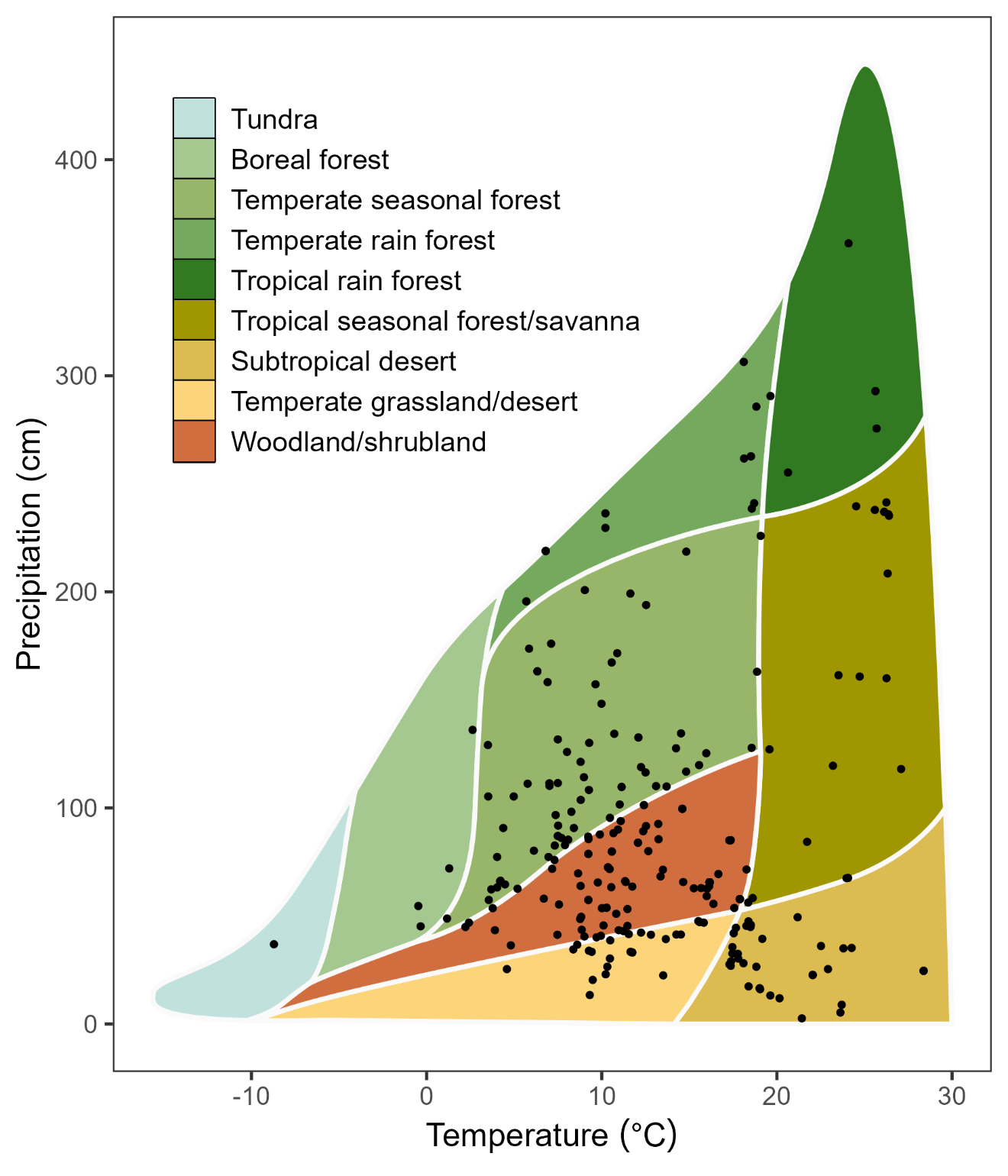

*Figure S2. Distribution of PSInet field studies (n = 272) in Whittaker biome space; mean annual temperature and mean annual precipitation extracted from WorldClim 1970-2000 normals.*

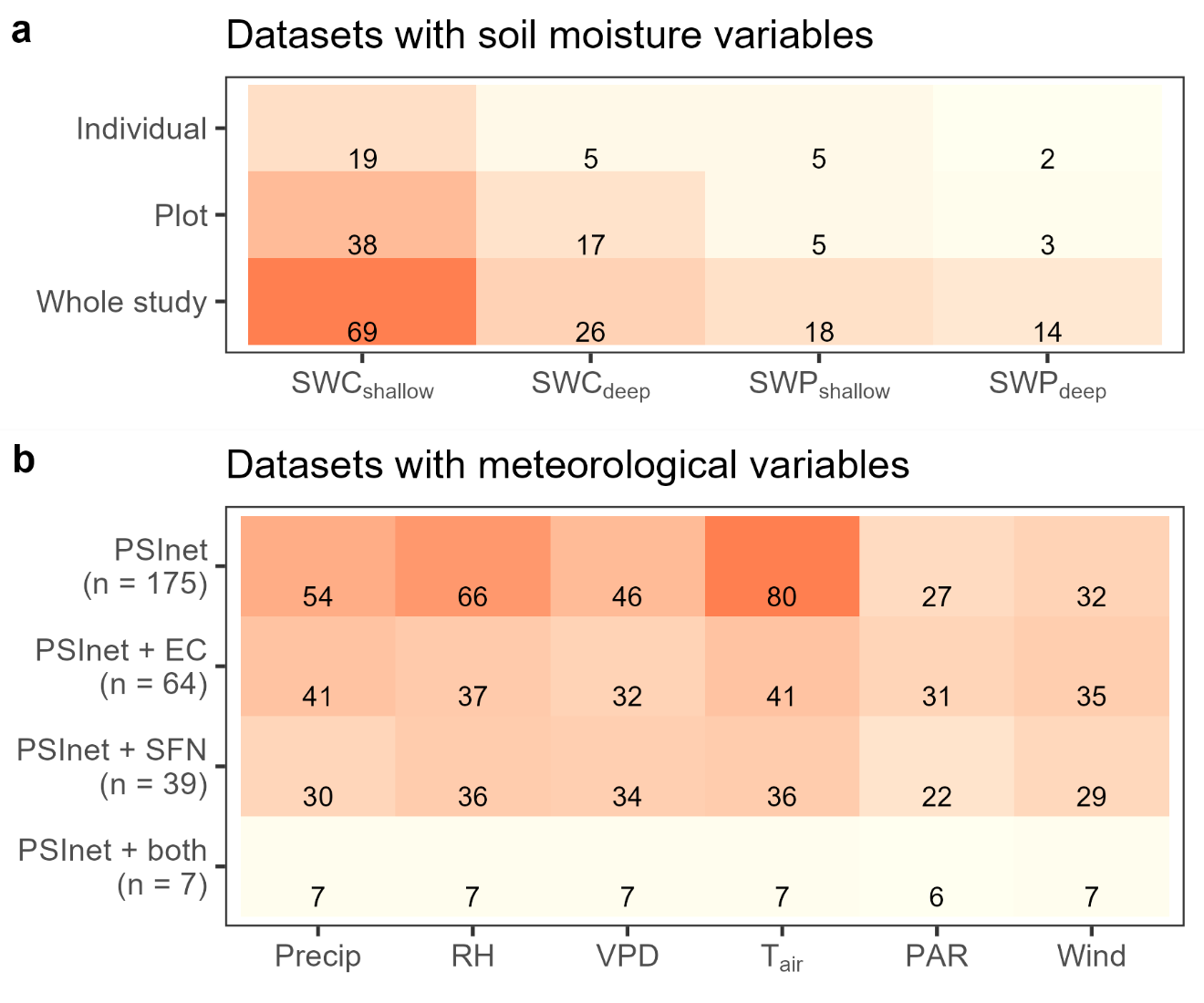

*Figure S3. Heatmap depicting the number of datasets for which (a) soil moisture variables were available by the level of organization (individual, plot, or whole study) and (b) meteorological variables were available by datasets from PSInet alone or co-located with eddy covariance towers (EC), SAPFLUXNET data (SFN), or both.*

*Table S1. Time series variables and their associated table, standard unit, and range. Values outside this range were marked as FALSE in the flag table but not removed from the database.*

| Table | Variable | Unit | Minimum | Maximum |
| --- | --- | --- | --- | --- |
| chamber_wp | water_potential_mean | MPa | -15 | 0 |
| chamber_wp | water_potential_sd | MPa | 0 | 1.5 |
| auto_wp | water_potential_mean | MPa | -15 | 0 |
| auto_wp | water_potential_sd | MPa | 0 | 1.5 |
| soil_var | swc_mean_shallow | g/g or cm^3^/cm^3^ | 0 | 1 |
| soil_var | swc_mean_deep | g/g or cm^3^/cm^3^ | 0 | 1 |
| soil_var | swp_mean_shallow | MPa | -50 | 0 |
| soil_var | swp_mean_deep | MPa | -50 | 0 |
| met_var | precipitation_mm | mm | 0 | 250 |
| met_var | relative_humidity_percent | % | 1 | 100 |
| met_var | vapor_pressure_deficit_k_pa | kPa | 0 | 12 |
| met_var | air_temperature_c | C | -30 | 50 |
| met_var | photosynthetically_active_radiation_ppfd_mumol_photon_m_2_s_1 | µmol photon m^-2^ s^-1^ | 0 | 2400 |
| met_var | incident_shortwave_radiation_W_m_2 | W m^-2^ | 0 | 1362 |
| met_var | net_radiation_W_m_2 | W m^-2^ | -300 | 2000 |
| met | windspeed_m_s | m/s | 0 | 45 |

**Supplementary Table 2:** *Describes key features of each contributed dataset, including dataset ID, the reference doi, the water potential measurement type, the study location and duration, and details on the number of species and individuals monitored. It is* [***available here***](https://drive.google.com/file/d/1sr7LItpIcy5mU28rgLxvXx9K7S3iqOPO/view?usp=sharing)

**Supplementary Table 3:** *Provides details on the availability of ancillary data for each PSInet dataset, including the availability of soil water and meteorological measurements, whether each site is part of SAPFLUXNET or a flux tower network, and the existing of water retention curves, leaf-level gas exchange data, and other plant hydraulic data from each study. It is* [***available here***](https://drive.google.com/file/d/1BduLCR3GBs1GsU_8KYBhVVlMpXIACBUq/view?usp=sharing)
